# The LacI hinge region balances binding stability against inducibility

**DOI:** 10.1101/2025.05.07.652604

**Authors:** J. Yuan, M. Lüking, S. Zikrin, BC Sen, E. Marklund, D. van der Spoel, D. Fange, J. Elf

## Abstract

Transcription factors (TFs) efficiently locate their target DNA sequences by combining three-dimensional diffusion and one-dimensional sliding on nonspecific DNA. To balance rapid sliding with strong specific binding, TFs were proposed to switch between search and recognition conformations. For *E. coli lac* repressor (LacI), the folding of the hinge helices has been implicated in the conformational switch. Here, we tested how mutations in the hinge region impact the search speed and binding stability. Based on molecular dynamics simulations, we selected two LacI mutants favoring either search or recognition conformation. We measured the binding kinetics of the mutants both *in vitro* on DNA microarrays with 2,479 different Lac operators and *in vivo* via single-molecule experiments. We conclude that a hinge region mutation causing less helix propensity enhances the specificity but reduces binding strength globally, while a hinge region mutation causing higher helix propensity has opposite effects. However, altered specificity impacts the search time less than expected. Instead, the major effect was impaired dissociation in response to IPTG induction for the strongly binding mutant. Together with earlier reports of affinity–inducibility trade-offs in LacI, our data support the model in which the hinge region governs a trade-off between binding stability and inducibility rather than between speed and binding stability.

## INTRODUCTION

Transcription factors (TFs) modulate gene expression in response to environmental and developmental cues(1, 2). They accurately and quickly bind to specific DNA sequences within a large excess of non-specific DNA sequences in the genome. For example, the *lac* repressor (LacI) of *Escherichia coli* (*E. coli*) can find its specific target sequence among the 4.6 million genome base pairs in approximately 3.5 minutes(3). Fast TF search is achieved by facilitated diffusion(4), where the TF combines 3D diffusion with 1D sliding along DNA, effectively extending the target size for diffusion-limited association to the TF sliding distance(5, 6).

In the sliding state, the TF should only interact weakly (<1.5 k_B_T) with the DNA, or else the protein would be trapped at non-specific sites(7). At the same time, the TF should bind the specific sites strongly (>15 k_B_T), or else they will not compete well with the overwhelming number of non-specific sites. Given a model where the TF has a static conformation, where interactions with different bases are additive, and there is a continuous distribution of binding energies, the combination of fast sliding and strong binding is not possible(7). This is known as the speed-stability paradox. A possible resolution to the paradox was suggested by Berg *et al*.(8). If the TF adopts two conformations, one for non-specific interaction with DNA (search conformation), and one for specific interaction with DNA (recognition conformation), and switches between the conformations either rapidly or guided by the sequence, the speed-stability paradox can be resolved(9).

The conformation-switching model is supported by the fact that several TFs have different conformations, which are often stabilized by interactions with small molecules and other proteins. This is the case for the LacI/GalR family in bacteria, which depends on the interaction with small molecular allosteric factors(10), as well as for some human transcription factors, such as the ones in the Myc family, which depend on dimerization for specific DNA-binding(11). The ability of TFs to adjust their structural and functional states in response to changes in the cellular concentrations of other molecules is crucial for their efficiency as genetic regulators(12). Often, intrinsically disordered regions play a key role in conformational switching at specific sites for transcriptional control and signalling(13).

The question remains whether favoring the search or the recognition conformation of TFs produces predictable changes in DNA specificity, binding stability, and target search speed within living cells. Specifically, it is unclear if evolution fine-tuned the TF’s balance between the search or recognition conformation to solve the speed-stability paradox, or if there are other physical constraints that are more important. For example it has been shown that increased binding strength to the operator may come at the cost of reduced inducibility(14). Here, we address the question for the *E. coli* transcription factor LacI using a combination of kinetics measurements *in vitro* and *in vivo* for *in silico-*designed LacI mutants that favor the hypothesised search or recognition conformation.

We specifically ask if (*i*) a preference for the search conformation comes at the expense of lower binding probability and reduced binding strength to its operator DNA, and (*ii*) if a preference for the recognition conformation causes strong interactions with non-operator DNA, making the search for the real operators prohibitively slow.

*Box*: Throughout the paper, binding strength, specificity, operator access time, search time, and association rate are defined as follows: *(i)* **binding strength** or affinity refers to the equilibrium binding constant for a DNA sequence; *(ii)* **specificity** refers to the difference in binding strength between an operator sequence and non-operator sequences. A reduction in specificity means reduction in the fold change difference between operator and non-operator sequences; *(iii)* **Operator access time** refers to the time for one LacI to reach the operator sequence in the cell but not necessarily binding to it. This time includes the time spent diffusing in 3D, binding and sliding on non-operator sequences that occurs before the operator is reached; (*iv*) the **search time** is the time it takes for the lac operator to be bound by a LacI. (*v*) The **association rate** is the inverse of the search time. The inverse relationship relies on the fact that searching for the stretch of DNA that contains the operator is a random non-sequential process and that the time spent non-specifically bound to the stretch of DNA that contains the operator is negligible in relation to reaching this stretch of DNA.

## MATERIAL AND METHODS

### Molecular Dynamic Simulations

An apo LacI dimer structure was constructed by removing DNA and ONPF from the 1EFA (15) crystal structure. Mutations to this structure were introduced using PyMOL(16) where the rotamer configuration resulting in the smallest amount of sterical clashes for each mutant were selected. The starting structures for the simulations were either the mutated (V52A, Q55N, or G58A) or the non-mutated (WT) apo LacI dimer described above. The simulations were performed with GROMACS version 2024.3(17) and the AMBER99SB-ws(18) force field. Water molecules were modeled using TIP4P/2005 water(19). Potassium parameters from Luo & Roux(20) were used, as standard AMBER force fields tend to overestimate the strength of non-bonded interactions for potassium(21). We ran in the NPT ensemble at a temperature of 310 K and in a solution that contains 150 mM KCl and 5 mM magnesium ions, based on previous simulations (22).The LINCS algorithm(23) was employed to restrain intramolecular bonds involving hydrogens. We used a 2-fs-time step for integration. Short-range interactions were calculated within 1 nm from the solute, while electrostatic interactions were calculated using the Particle-Mesh-Ewald method(24). The stochastic velocity rescaling thermostat (25) and C-rescale barostat(26) were used for temperature and pressure coupling, respectively. In preliminary simulations, the DNA binding domain (DBD) was observed to stick to the core domain in a manner which we deemed incompatible with non-specific binding. To ensure that this conformation of the DBD and the core domain did not bias our helix propensity estimates, distance restraints were introduced between the DNA binding domain and the core. Here flat bottomed harmonic potentials were used. Alpha carbons on the following residues were used to add restraints: T34, E39 and D43 on the DBD, D88 on core domain on the same monomer as the selected DBD residues, A106 and A110 on the core domain of the opposite monomer in the LacI dimer. Lower bounds between 1.3 and 2.2nms, were added to the following pairs T34-A106, T34-A110, T34-D88, E34-A106, E34-A110, E34-D88, A43-A106 and A43-A110. See Table S6 for details. Upper bounds were set to 4.2nm for all pairs. The lower and upper bounds were set heuristically using simulations without distance restraints. The lower and upper bounds were set heuristically based on simulations without distance restraints. RMS deviations from an ideal helix (Fig. 1E) were calculated for residues Arg51-Ala57 using the Gromacs helix function. For each time-point the hinge region was classified as helical if the RMSD<0.1nm. The fraction of helical hinge regions were calculated for each time point and for each hinge region assuming that the two hinges in the LacI dimer fold and unfold independently.

**Figure 1:**
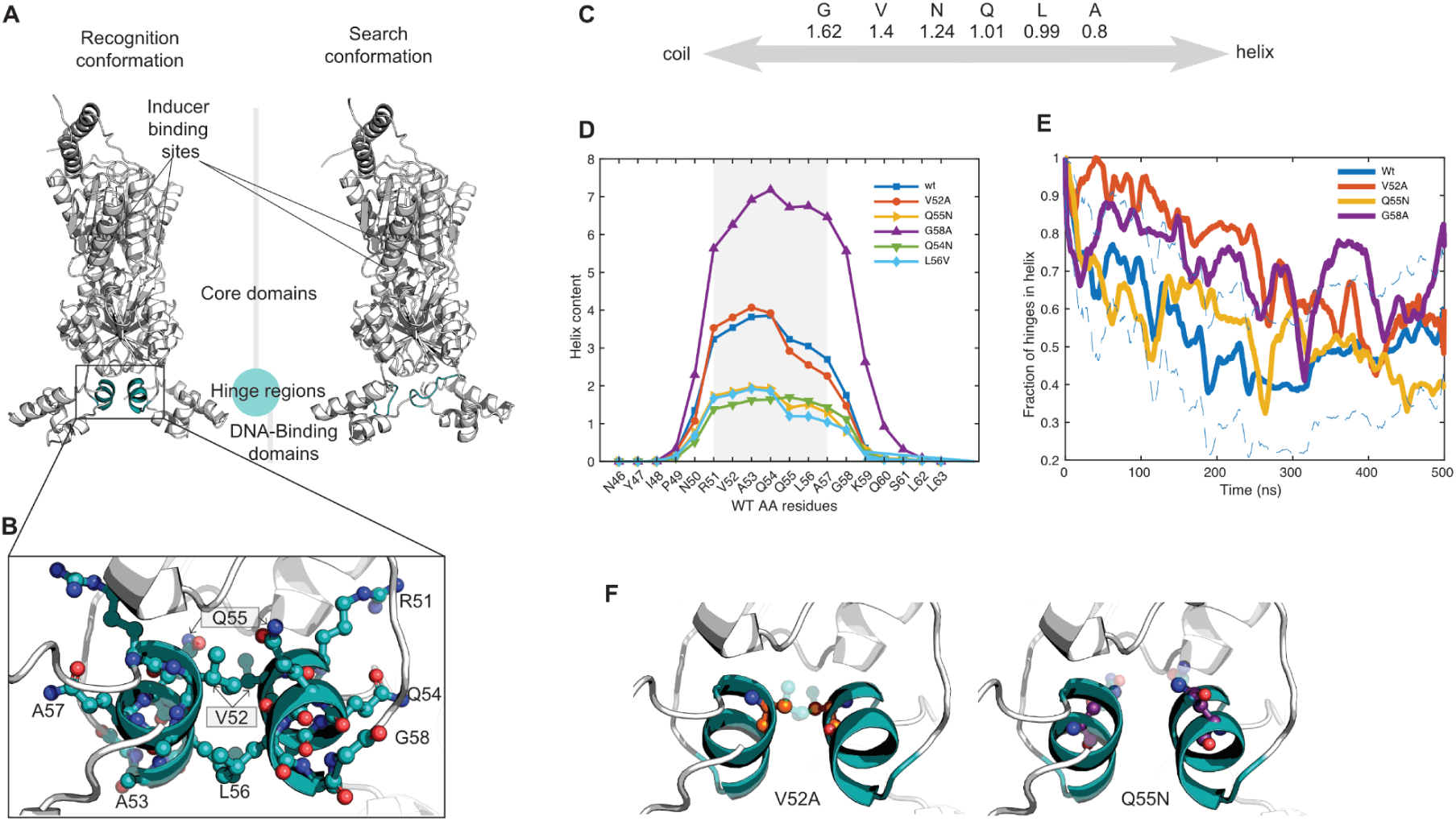
In silico analysis of point mutations in the LacI hinge region regarding their impact on helix propensity. **A.** Cartoon representation of LacI dimer conformations. (Left) Recognition conformation (PDB ID: 1EFA) with hinge region highlighted in teal. (Right) Hypothetical search conformation visulized by integrating the core domain from 1EFA with the DNA-binding domain from a NMR structure (PDB ID: 1OSL, (41)), as described in Lüking et al.(42), with hinge region similarly highlighted in teal. **B.** Cartoon and Ball-and-Stick representation of the hinge helix residues. In the Ball-and-Stick model: Carbon atoms are shown in teal, nitrogen atoms in dark blue, and oxygen atoms in red. Residue names are labeled. **C**. Subset of estimated characteristic energies for helix folding (excluding the formation of hydrogen bonds) in kcal mol^-1^ of different protein residues as presented by Muñoz and Serrano(43). **D.** Helix content per residue calculated by Agadir (ref) for 18 amino acids centered around part of the hinge which is in helical form in the 1efa crystal structure (gray). **E.** MD simulation outcomes. Fraction of hinges which have helical form (RMSD<0.1nm in Fig. S3) as a function of time (solid lines). The averages have been smoothed using sliding window average with window size 10ns. The dashed lines show 1SEM around the smoothed mean for each time point. The SEM has been smoothed using a sliding window average with window size 10ns. The RMSD from ideal helix for residues R51-A57 for each of the five simulations each containing two hinges (LacI dimer) are shown in Fig. S3. **F.** Cartoon and Ball-and-Stick representations of the point-mutated residues (V52A in orange and Q55N in purple) introduced into the hinge region of Wt-LacI (See point-mutation methods detailed in Methods). The original wild-type residues (shown half-transparent) are included for comparison and correspond to those in panel B.

### Construction, expression, purification, and fluorophore labeling of LacI variants

The LacI-Halo constructs used in the *in vitro* DNA microarray experiments have the same LacI sequence as in our previous study (27). Additionally, a HaloTag (underlined in sequence info in Table S3) and a 6xHis-Tag were translationally fused to the C-terminal of LacI. LacI mutants were constructed based on the LacI-Halo construct. The V52A-LacI-Halo construct introduced an Alanine at amino acid position 52 of the LacI-Halo protein to replace Valine. The Q55N-LacI-Halo construct introduced an Asparagine at position 55 to replace the original Glutamine. See sequences of LacI-Halo constructs in Table S3.

All variants of LacI-Halo were expressed in BL21 *E.coli* cells using the pD861 plasmid, which is taken from Marklund *et al.*(*28*). The resulting strains—EL4464, EL4465, and EL4466—were used for overexpression of the Wt, V52A, and Q55N LacI variants, respectively. The protein expression and purification protocol, adapted from a previously published method, is described below (28). 5 ml overnight cultures were diluted into 500 ml LB medium with 50 µg/ml kanamycin and grown at 37 °C and 110 RPM agitation. The expression of LacI was induced at OD600 = 0.5-1 by adding L-rhamnose to a final concentration of 0.2 % (weight/volume). After 3 hours, the cells were harvested by centrifugation at 6,000 x g at room temperature for 40 min. The pellet cells were then resuspended in 25 mL BugBuster® Master Mix (Merck) solution containing 1 EDTA-free protease inhibitor pill (Roche), and the cell suspension was incubated on a shaking platform at 50-75 rpm for 10–20 min at room temperature. The lysate was then clarified by centrifugation at 5,000 x g for 1 hour at 4 °C, followed by filtering of the supernatant through a 0.45 µm cellulose acetate membrane filter. The collected soluble extract was loaded directly onto a His GraviTrap column (Cytiva). The LacI-containing fractions were pooled and buffer-exchanged three times using 50 kDa cut-off Amicon Ultra-15 Centrifugal filters (Merck Millipore) into phosphate-buffered saline (PBS). Samples in PBS were diluted by adding an equal volume of 90% glycerol in PBS, resulting in a final storage solution of 45% glycerol in PBS. The protein was then kept at –20 °C for short-term storage prior to labeling.

The labeling of LacI proteins with HaloTag® TMR Ligand (Promega) was performed in PBS + 5% Glycerol. A volume of purified protein stored in 45% Glycerol in PBS, as previously described, was mixed with 9 volumes of PBS, and an excess amount of TMR ligand (at least a protein:ligand ratio of 1:1.1) was added to the labeling mixture. The labeling reaction mixture was incubated for 1 hour at room temperature. The labeled protein was loaded onto a His GraviTrap column (Cytiva) with binding buffer (20 mM Sodium Phosphate, 500 mM NaCl, 20 mM Imidazole), and the eluted fractions were collected using the elution buffer (20 mM Sodium Phosphate, 500 mM NaCl, 500 mM Imidazole). The eluate was then buffer-exchanged into 1× PBS using an Amicon® Ultra Centrifugal Filter (50 kDa molecular weight cutoff). Protein concentration and labeling efficiency were determined for the TMR-labeled LacI variants using absorbance measurements at 280 nm (A₂₈₀) and 548 nm (A₅₄₈). The protein concentration was calculated as in Thermo Fisher Tech paper(29): Protein concentration (M) = (A₂₈₀ – (A₅₄₈ × CF)) / ε, where ε is the protein molar extinction coefficient (81360 M⁻¹cm⁻¹), CF (correction factor) was measured as 0.23. Labeling efficiency, expressed as moles of dye per mole of protein, was determined using: Moles dye per mole protein = A₅₄₈ / (ε’ × protein concentration (M)), where ε’ is the molar extinction coefficient of the TMR ligand (78000 M⁻¹cm⁻¹). Absorbance values were averaged across at least five technical replicates. Proteins in PBS were then diluted to the desired concentration with an equal volume of 90% Glycerol in PBS and were aliquoted for long-term storage at –80 °C.

### Microscopy of protein binding microarrays

The design and protocols for PBMs and microscopy were replicated from a prior study(27) with the addition that the LacI variants could be flowed into different PBM-containing flow chambers on the same device (Fig. S7), and thus, the kinetics of different variants could be measured simultaneously.

To prepare the single-stranded oligonucleotide microarrays synthesized by Agilent Technologies as PBMs, they were first synthesized into double-stranded DNA using a reaction mixture containing Cy5-dCTP according to previously published methods (27, 30). Prior to the association and dissociation experiments, PBMs were incubated with imaging buffer (Potassium phosphate 10mM, EDTA 0.1mM, Glycerol 5%, Sodium Chloride 20mM, 1mM 2-Mercaptoethanol, 0.5mg/ml BSA, 2% non-fat milk (Semper), final pH were adjusted with 1M Potassium hydroxide to 7.2) for 1 hour at room temperature (∼21°C). After incubation, association experiments were started by flowing 1.85 nM LacI dimer and 1.85 nM mutant-LacI dimer in Imaging Buffer into different flow cells at 450 µl/min. Four to five hours after the start of the association experiments, the dissociation experiments were started by flowing in the imaging buffer without LacI proteins at 1 ml/min.

#### Microscopy and Imaging

The microscope setup is the same as described in our previous study (27). Two flow cells were imaged together in one round of imaging, each location on these two PBMs was imaged every 51 seconds through the TMR channel for the first 60 time-points of association experiments, followed by 5-minute intervals thereafter. In dissociation experiments, imaging was conducted every 51 seconds for the first 36 scans, then shifted to 5-minute intervals.

### Analysis of reaction kinetics from PBM measurements

#### Image Analysis

This image analysis workflow is designed to accurately quantify TMR fluorescence intensities from PBM images, while addressing spatial heterogeneities observed in binding patterns. These patterns are shown as elevated fluorescence at the front-flow-facing region of DNA spots, especially DNA spots that show strong affinity to the flowed LacI variant. We hypothesized that this effect resulted from the depletion of free LacI downstream of the flow front near each spot. To accurately quantify spot binding intensities unaffected by this phenomenon, we refined our previously established image analysis method(27) as described below.

Initially, following our earlier approach, merged flow-cell frames—each consisting of around 40 tiled images—were stitched together. The initial binary masks for DNA spots (*maskIni*) and their local backgrounds (*bkgMaskIni*) for an array in one PBM experiment were generated using known spot centroids and derived spot radius (*spotRadius = rowSpacing / 3.9*) from Agilent’s microarray geometry specifications, which provides information including *rowSpacing*. For each DNA spot, its initial fluorescence intensity was computed by the difference between its mean spot intensity value and the corresponding local background mean intensity. Invalid spots, which are located within damaged regions (e.g., scratches, air bubbles, or tubing coverage), were manually masked using Fiji software and were excluded from further analysis.

Next, to refine spot segmentation further, DNA spots exhibiting higher initial fluorescence intensity than those corresponding to the *O_1_* operator in the first dissociation experiment frame (*im1stDissoc*) were selected. These high-affinity spots were selected because, first, they are prone to show inhomogeneous binding during early association and, secondly, they can be used to improve the localization of DNA spots. A more precise spot localization was needed because accessing the flow-front-facing region of spots requires a precise on-edge localization of whole DNA spots.

Pixels in the *im1stDissoc* image that are farther than twice the spot radius from high-affinity spot centers were replaced with local median background values, creating a refined image. Bright circular spots in this refined image were detected using MATLAB’s function imfindcircles. An affine transformation matrix that aligns the centroids of these spots with the paired spots’ centroids in *maskIni* was computed using MATLAB’s function fitgeotrans. The *maskIni* was updated, as *maskTemp*, by moving all spot centroids as specified by the inverse of the transformation matrix and expanding them slightly (1.1 × spotRadius). A separate background mask (*bkgMask*) was similarly transformed.

High-affinity DNA spots, now accurately segmented with *maskTemp*, were then further processed for identifying the top 20% fluorescence intensity regions within each spot at an early association time point (t=1,650 s), where the inhomogeneous binding pattern, by visual inspection, were evaluated to be the most prominent. The spot segmentation for these high-affinity DNA spots was then replaced with their identified top-20%-intensity regions in *maskTemp*. Weak-affinity spots inherited these 20% intensity regions from their nearest neighboring high-affinity spots. Together with the replaced spot segmentation for high-affinity spots, we have a final binary mask for DNA spot segmentation addressing the heterogeneous binding pattern, *maskFinal*.

The final spot fluorescence intensity time-trace for each DNA spot was computed as the difference between its mean intensity value from the top 20% segmented region segmented by *maskFinal* and their corresponding local background mapped by *bkgMask*. These refined segmentation masks for DNA spots and local background (*maskFinal* and *bkgMask*) were uniformly applied to all image frames in one PBM experiment with a certain LacI variant; all frames were first subtracted with an offset frame captured before the introduction of LacI into the flow cell for the purpose of minimizing background noise. Consequently, the time-resolved fluorescence intensity trace for each valid DNA spot across association and dissociation phases in a PBM experiment involving one LacI variant was then obtained.

#### Post-processing and Kinetic Analysis of Acquired PBM fluorescence time traces

Obtained DNA spot fluorescence time traces were first filtered and then grouped by corresponding DNA sequence. Spots were filtered according to four criteria: *(i)* Each spot’s initial local background must not exceed 400 fluorescence units (the background level during the association phase). This threshold ensures the exclusion of DNA spots that cannot be identified for their exact time points for the beginning of association. *(ii)* The maximum value, *f_max_*, of moving means (10 time point window size) of the spot intensity time trace needs to be > 0. *(iii)* The sum of the absolute differences between subsequent intensity values in the time-trace needs to be below a threshold, 85, because large frame-to-frame variability in the time trace is not consistent with LacI binding to DNA. (*iv*) Each DNA spot’s local background time trace was normalized to its *f_max_*, and only those normalized traces with fewer than six values falling outside of the range of –0.5 to 1.5 were kept. The spots that passed all four filters above were assigned to their corresponding unique DNA sequence group. DNA sequences that already had < 3 valid spots from this initial filtering step were discarded from further analysis.

Next, the exact time points marking the beginning of association and dissociation were determined for each valid DNA spot. When the reaction buffer containing fluorescently labeled LacI protein flowed into the PBM chamber, the background fluorescence increased rapidly due to the presence of free protein in the Imaging buffer. Conversely, when the LacI-containing buffer was replaced by the Imaging buffer without protein, the background fluorescence rapidly decreased. To identify precisely when these buffer exchanges occurred for each DNA spot, we first obtained rough estimates using the average overall background intensity across all DNA spots on the whole array. Specifically, the time point where the largest increase happens in the overall background intensity time trace indicates the approximate universal start of the association phase, and the time point with the largest decrease indicates the universal start of the dissociation phase for all DNA spots. Then, for each individual DNA spot on this array, we refined these rough start time estimates. The average high-local-background fluorescence during the association phase of each DNA spot was computed by averaging the intensities from 10 consecutive time points, beginning 10 time points after the estimated universal start of association time point in its local background time-trace. Similarly, the low-local-background intensity during its dissociation phase was calculated. Each DNA spot’s local background fluorescence time trace, after being subtracted with the low-local-background value, was then normalized to its absolute difference between its own high– and low-local-background values. Then, the exact start time point index of association for each DNA spot was defined as its first time point index, *startIdx*, where the normalized local background fluorescence value exceeded 0.5. Similarly, the dissociation start was determined as the last time point index, *stopIdx*, meeting this criterion. After identifying the start and stop indices, DNA spots with unrealistic indices were excluded from analysis based on the following three criteria: *(i)* association to dissociation duration fewer than 50 time-points recorded, *(ii) stopIdx* is beyond the selected range of corresponding time-frame with < *bkgLow* intensity, *(iii) startIdx*=1, because this means the association on this DNA spot already started before we could capture it. For valid DNA spots that pass the above criteria, the precise association and dissociation times (*startTime* and *stopTime*) were then interpolated between their corresponding time points for frame index: *startIdx*-1 and *startIdx*, *stopIdx*-1 and *stopIdx*, respectively.

The binding and unbinding kinetics for each DNA spot were modeled by fitting their fluorescence time traces to exponential models. In the association phase, a single exponential curve is applied to fit the intensity data, *f(t)*, over a 30-minute time window from the interpolated *startTime*. The association fit function was defined as 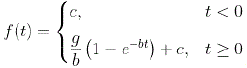, where *g = k_a_\** [Protein]*[DNA_tot_]*proteinFluor, where [Protein] means the free concentration of LacI during the flow experiments, [DNAtot] means the total concentration of double stranded DNA on a DNA spot and proteinFluor means the mean fluorescence each protein gave, *b* = *k_a_*\* [Protein] + *k_d_*, and c is a constant. For the dissociation phase, a 15-minutes window after the interpolated *stopTime* was used.

An inferred equilibrium fluorescence, *C_max_*, was calculated as 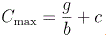. During the first 15 minutes of dissociation, the DNA spot intensity curves are well-described by the sum of an exponential decrease and a constant. To estimate the dissociation rate constant, *k_d_*, we use the initial slope of the fitted curves. More specifically, two dissociation fitting models were employed for weak and strong affinity DNA spots, respectively. For weak-affinity DNA spots (those with computed maximum value out of their 10-frame moving means from their time trace < 150 fluorescence units), 50 minutes prior to the dissociation start time were also added to the fitting window for better capturing the spot intensity at the dissociation start point. In this first scenario, the expanded fitting window used the function:

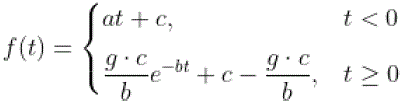

Here, *c* is the spot intensity at the start of the dissociation. The initial slope of the dissociation curve, *f’*(*0*) *= – g * c*, and thus *g = k_d_*. For spots with higher intensities (≥150 fluorescence units), the constant *c* is fitted directly from the dissociation curve, and thus a simpler model was used: 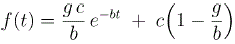, where *g = k_d_.* The underlying reason for the constant offset is unknown.

To assess the reliability of the fitted *g* parameters from both the association and dissociation stages for a DNA spot intensity time trace, we calculated the coefficient of variation (CV) by dividing the half width of the 67% confidence interval of the *g* parameter (approximately one standard deviation) by the corresponding g value for both fits. DNA spots for which the CV > 1 for either the association or the dissociation stage—indicating that the relative uncertainty was greater than the fitted parameter itself—were excluded from further analysis.

For each unique DNA sequence represented by at least 3 valid DNA spots in a LacI variant-PBM experiment, its averaged ‘equilibrium’ fluorescence (*f_eq_*) and corresponding SEM were computed based on *f_max_* values from all its valid DNA spots. Finally, averaged kinetic parameters and their corresponding SEM were determined from fitted parameters as listed below for each valid DNA spot: the association parameter *g*, equal to *k_a_*×[Protein]×[DNA tot]×proteinFluor; the inferred equilibrium fluorescence (*C_max_*); and the dissociation rate constant *k_d_*.

For data presented as in Figure 2 and S1, the kinetics parameters for each LacI variant with a unique DNA sequence, its association-related parameter g = *k_a_*\**C*, *C* = [Protein]×[DNA_tot_]×proteinFluor collects global experimental factors. All experiments were run with a fixed concentration of fluorescently labelled protein. In each replicate with two PBMs, the PBMs with printed ssDNA were synthesized into dsDNA in the same reaction, thus the [DNA_tot_] of all DNA spots on PBMs were assumed same within each replicate (this assumption is validated because no consistent spot-wise correlation were shown in one array for DNA spots of *O_sym_*,, *O_1_* or *O_2_*, data not shown). Taken together, *C* cancels when we normalize *k_a_*\**C* with Wt-*k_a_*(*O_sym_*)\**C* within a side-by-side PBM experimental replicate. These within replicate relative *k_a_*values were then averaged across replicates to obtain the reported relative *k_a_* values.

**Figure 2.**
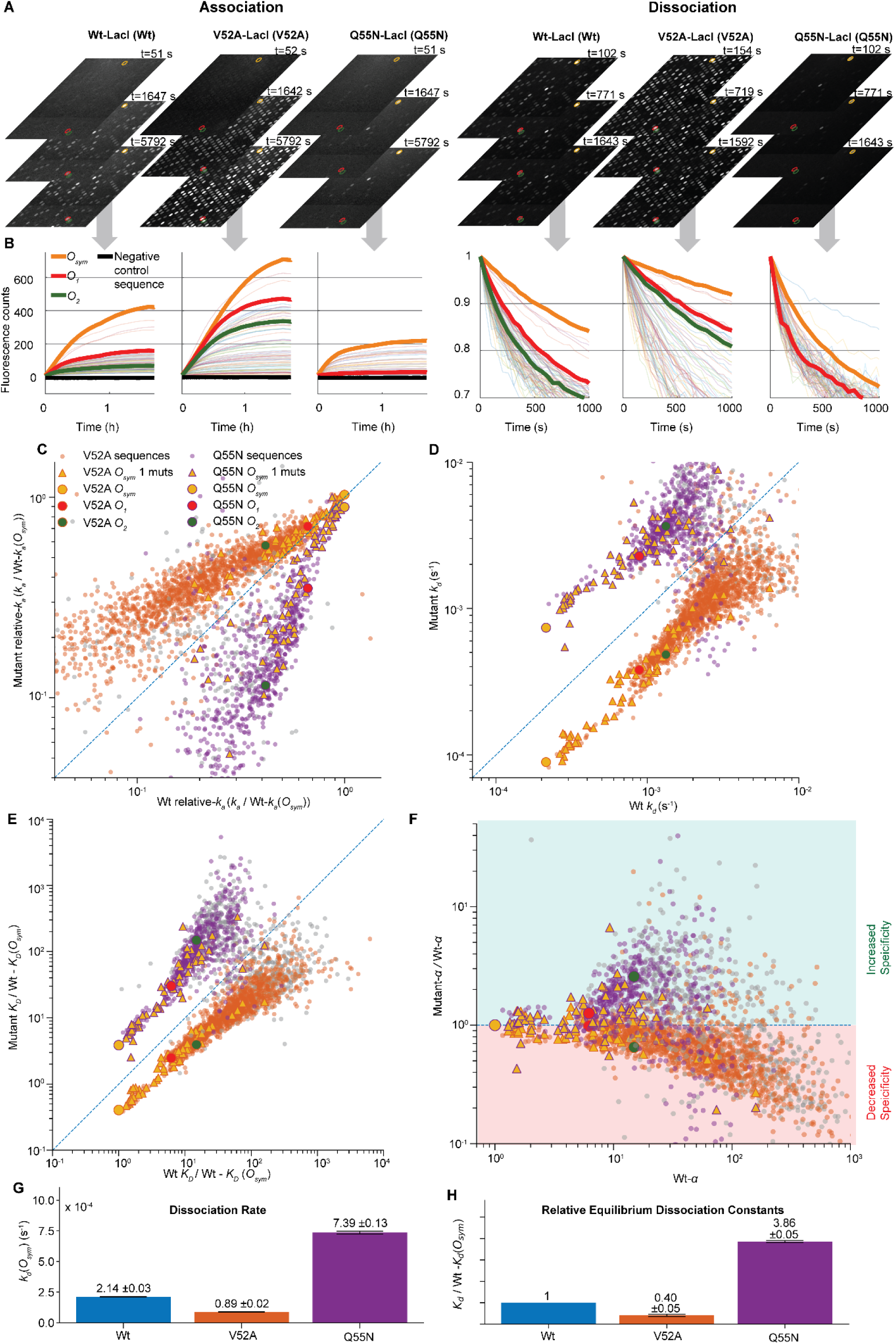
*In vitro* real-time kinetics measurement for LacI variants on PBMs. **A**. Representative fluorescence time-lapse images of association (*left*) and dissociation (*right*) phases for Wt-LacI (Wt) and mutant LacI variants (V52A and Q55N) on the DNA microarrays. **B**. Real-time binding curves for *O_sym_*, *O_1_*, and *O_2_* operators and random 80 examples of their mutants (faint color), along with the negative control sequence. The association curves (*left*) show the increase of fluorescence intensity over time, while the dissociation curves (*right*) display the normalized fluorescence decrease from the first frame of the corresponding dissociation movie. Each curve corresponds to fluorescence intensity changes over time for a unique DNA sequence within a single PBM experiment using a specific LacI variant. The fluorescence values shown represent averaged intensities from at least three replicate DNA spots within that single PBM experiment. The curves presented here have been filtered to exclude any DNA sequences whose fitted association (*k_a_*) or dissociation (*k_d_*) rate constants showed a coefficient of variation (CV) greater than 1; these sequences were also omitted from the calculations of the mean relative-*k_a_* and *k_d_*values displayed in panels C&D. **C**. Correlation of relative-*k_a_*values (normalized with Wt-LacI *k_a_*(*O_sym_*) value) for unique DNA sequences between Wt-LacI (x-axis) and LacI mutants (y-axis), with V52A (orange) and Q55N (purple) LacI variants plotted in the same panel. Sequences with high variability in relative-*k_a_*measurements (CV > 0.3) across corresponding replicates are marked in grey. Each data point represents the mean relative-*k_a_* value for a unique DNA sequence, calculated from all binding spots with the same sequence in 10 replicated experiments for Wt-LacI and 5 replicates for each mutant-LacI (See individual side-by-side Wt vs Mutant PBM experiment in Fig. S5A). **D**. Correlation of dissociation rates, *k_d_*, between Wt (x-axis) and its mutants (y-axis), Sequences with high variability in *k_d_* measurements (CV > 0.3) are marked in grey (see individual side-by-side Wt vs Mutant PBM experiment in Fig. S5B). **E.** Correlation of equilibrium dissociation constants (*K_D_*) between Wt (x-axis) and LacI mutants (y-axis). *K_D_* values, calculated from *k_d_* / *k*_a_, of each protein to different sequences are normalized with the *K_D_* value of Wt to the *O_sym_* operator (see individual side-by-side Wt vs Mutant PBM experiment in Fig. S5C, and Fig. S5D for *K_D_* derived from 1/ f_eq_, f_eq:_ fluorescence measured at / close to equilibrium). **F.** Specificity changes were assessed using dissociation constant (*K_D_*) estimates. For each DNA sequence (i), a LacI variant’s *K_D_* to it was defined as the product of its *K_D_* for the *O_sym_* and a specificity factor, *α*(i), *i.e. K_D_*(i)=*α*(i) *K_D_*(*O_sym_*). If a mutation in LacI does not alter its DNA specificity to *O_sym_*, the *α* values for Wt-LacI and Mutant-LacI would be identical for all sequences, *i.e* Wt-*α*(i) = Mutant-*α*(i) for all *i*. Normalized Mutant-*α* values (relative to Wt-*α* values) were plotted against Wt-*α* values. If a LacI mutant’s normalized α values are generally bigger than 1 (green area), that means, compared to Wt, its specificity to *O_sym_* is increased, while values below 1 (red area) indicate that its specificity to *O_sym_* is decreased. **G.** Measured *k_d_*(*O_sym_*) values, shown as mean ± SEM **H.** *K_D_*(*O_sym_*) values are calculated from *k_d_*/ *k_a_* values and normalized with Wt-*K_D_*(*O_sym_*), shown as mean ± SEM.

#### Calculation for Microscopic Rates

As shown in Marklund *et al.* (27), the microscopic dissociation rate (*k_off,µ_*) and the maximum association rate constant (*k_on,max_*) can be inferred by applying the equation, 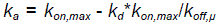. We fitted the above equation to the observed *k_a_*, *k_d_*relations shown in Figure S1A. The *k_on,max_* value and its standard error (SE) were computed through fitting the linear regression on *O_sym_* single mutant sequences showing the top 35% *k_a_* among the sequences that have their *k_a_*and *k_d_* both satisfy their coefficient variation (CV) < 0.3. For estimating *k_off,μ_*, the above determined *k_on,max_* was used in the equation: 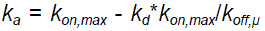, which was applied to all valid *O_sym_* single mutant sequences that have both *k_a_* and *k_d_* passed the CV < 0.3 filter. The mean and the standard error of the mean (SEM) of *k_off,μ_* were then calculated after trimming the top and bottom 10% of the individual calculated k_off,μ_ values to remove the outlier values.

### Strain construction for *in vivo* experiments

#### For microscopy (single-molecule experiment)

The background strain with the wild type LacI is *E. coli* BW25993 *lacI-mVenus* Δ(*p*_lac_*-lacZ*)::term, named EL3498(3). Using DIRex(31), point mutations were introduced in the *lacI* gene of EL3498 to obtain strains, EL3518 with LacI(V52A), and EL3520 with LacI(Q55N) amino acid substitutions. PCR was performed using primer pairs, aslA-hom-cat-*O_sym_*-Fwd and aslA-hom-cat-*O_sym_*-Rev (see Table S2) to obtain PCR product, lac*O_sym_* (sequences included in the reverse primer) along with Chloramphenicol cassette, herein written as, cat-lac*O_sym_*. As a template for Chloramphenicol resistance cassette, chromosomal DNA of strain EL8 was used. This PCR product was DpnI digested and further introduced into strain EL3498, using lambda red transformation(32) to obtain a strain with cat-*O_sym_* in the *aslA-glmZ* intergenic region. The resulting strain was named EL4053. P1 transduction(33) was performed from strain EL4053 to transduce cat-lac*O_sym_*into the recipient strains, EL3498, EL3518, and EL3520 to obtain strains, EL4089, EL4242, and EL4243, respectively.

As we aimed to have only a few LacI molecules per cell in order to keep the fluorescence from non-bound LacI low, we reduced the expression of *lacI* in the above strains by changing the wild-type *p_lacI_* promoter to the *p*FAB138 from the Mutalik *et al.* promoter library(34). A ribosome binding site (RBS from *p_lacI_* (wt)), was added after the promoter. A kanamycin resistance cassette was introduced upstream of the promoter sequence, transcribing in the opposite direction of the pFAB138 promoter. PCR primers were designed such that the primer, Comp1_Kan-Rv, has 40 bp homology at the *mhpA-mhpR* region followed by bases to amplify the kanamycin cassette from the reverse side (3’ end), while the other primer pair, Comp2_lacI 14 bp_RBS_p138-Fw contains bases to amplify the kanamycin resistance cassette from the forward side (5’ region), *p*FAB138, RBS and the first 14 bases of *lacI*. As a template for the kanamycin resistance cassette, chromosomal DNA of strain EL2620 was used. The obtained PCR product was DpnI-digested and introduced into EL3498, EL3518, and EL3520 by lambda-red transformation to yield the strains EL4210, EL4264, and EL4265, respectively. Thereafter, P1 transduction from EL4210 (donor) into EL3498 and EL4089 yielded the strains, EL4219 and EL4226, with and without cat-lac*O_sym_*, respectively. Similarly, P1 transduction from EL4264 (donor) into EL3518 and EL4242 yielded the strains EL4266 and EL4268, respectively. Furthermore, P1 transduction from strain EL4265 (donor) into EL3520 and EL4243, yielded the strains EL4267 and EL4269, respectively.

#### For Miller assay

Strains with LacI-mVenus variants were constructed based on the microscopy strains as described above. Using infusion cloning (Takara Bio), cat-*p*_lac_-lac*O_sym_*-*lacZ* was cloned on a R6K plasmid. Genomic DNA of EL409 was used as the source for *p*_lac_-lac*O_sym_*-*lacZ*. The resulting plasmid, pBS19, was sequence verified using primer pairs T7 Term_fwd and CAT-R. The insert of interest, herein written as cat-lac*O_sym_*-*lacZ,* contains the Chloramphenicol cassette upstream to the lac promoter sequence transcribing in the opposite direction, followed by lac*O_sym_* and the entire *lacZ* gene. PCR was performed using plasmid template pBS19 and the primer pairs aslA-hom-cat-*O_sym_*-F and cat-*O_sym_*-lacZ_aslA_R. The PCR product was DpnI-digested and introduced at the *aslA-glmZ* intergenic region of strain EL4219, EL4266, and EL4267 using lambda red transformation(32) to obtain the strains, EL449, EL4493, and EL4495, respectively. To obtain a *lacI* knock-out mutant as a background strain for the Miller assays, we first exchanged *lacI* in the strain EL4219 with a selectable/counterselectable marker, *Acatsac1*(*31*). Further, *Acatsac1* was replaced by a curing oligo tagatttaacgtataagagagtcaattcagggtggtgaatatggtgagcaagggcgaggagctgttcaccggggtggtgc, resulting in the strain EL4536. The PCR product, cat-lac*O_sym_*-*lacZ* from above, was further introduced at the *aslA-glmZ* intergenic region of EL4536 using lambda red transformation(32), resulting in the strain EL4580.

Strains for LacI-Halo variants, as used in DNA microarray experiments, were constructed similarly to above. We first exchanged the entire *mhpR-lacI-venus* in the strain EL3498 with the selectable/counterselectable marker *Acatsac1*(*31*). The resulting intermediate strain was named EL4621. Alongside, using infusion cloning (Takara Bio), Kan^R^-*p*FAB138-*lacI-halo* or Kan^R^-*p*FAB138-*lacI*(V52A)*-halo* or Kan^R^-*p*FAB138-*lacI*(Q55N)*-halo* was cloned on R6K plasmid, resulting in the plasmids pBS20 (Kan^R^-*p*FAB138-*lacI-halo*), pBS21 (Kan^R^-*p*FAB138-*lacI*(V52A)*-halo)* or pBS22 (Kan^R^-*p*FAB138-*lacI*(Q55N)*-halo*) respectively. All three plasmids were sequence verified using primer pairs Seq_Halo_F, Seq_Halo_F and lacI mid rev3252. PCR was performed using plasmid templates, pBS20, pBS21, or pBS22, and primer pairs Comp–Kan–R and LacI_Halo_R. PCR products were digested with DpnI and introduced using lambda red transformation(32) into the intermediate strain EL4621 and selected on sucrose plates. The resulting sucrose-resistant recombinants were named EL4630, EL4632, and EL4634. Finally, the PCR product, cat-lac*O_sym_*-*lacZ* from above, was transformed using lambda red transformation(32) into the strains, EL4630, EL4632, and EL4634, resulting in the strains, EL4639, EL4641, and EL4643, respectively.

All above strains used in single molecule experiment and Miller assay were sequence verified, however EL4643 showed a point mutation, 98C>98T (Pro34>Leu34), in the Halo-tag. We argue that this point mutation in halotag most likely does not influence the functionality of LacI in the Miller assay. Growth rate analyses were performed on the strains prepared for the Miller assays, all of which demonstrated comparable doubling times.

### Single-molecule tracking in living cells with microfluidics

#### Cell Culture and Preparation

Overnight cultures were grown in LB media at 37°C, diluted 1:500 in M9-Glucose media (0.4% glucose, 0.4% RPMI amino acids (Sigma-Aldrich R7131), and 80 μg/ml Pluronic F-109(Sigma-Aldrich 542342), and incubated for 2-3 hours at 30°C before loading into the microfluidic chip.

#### Microfluidic Chip and Media Flow Control

A PDMS microfluidic chip with cell trap-dimensions of 48 µm x 60 µm x 1 µm was used. Two different strains were loaded into opposite sides of the same microfluidic chip. In total 100 traps for each strain were imaged in each experiment. The microfluidic chip allowed for on-chip switching(35) between growth medium lacking IPTG to growth medium containing 0.3mM IPTG. Pressures in the chip was controlled using an in-house pressure regulator. Before the start of each association experiment, cells were exposed to 0.3 mM IPTG by switching pressures on the two media ports (from 90 to 240 mbar on the IPTG-containing media port and from 240 to 90 mbar on the IPTG-lacking media port) for at least 2 minutes.

#### Microscopy and Imaging

Wide-field microscopy was performed with the following setup:

● Microscope: Nikon Ti2-E with 100x/1.45 NA oil immersion lens (CFI Plan APO lambda, Nikon).
● Illumination: Spectra III (Lumencor) for epi-fluorescence.
● Optical components: Dichroic mirror (ZT514rdc-UF2 (Chroma)), excitation filter (FF01-514/3(Chroma)), emission filter (ET550/50M (Semrock)).
● Camera: Kinetix (Teledyne Photometrics).
● Software: micro-Manager (36).

Fluorescence images (4-second exposure) and phase contrast images (80 ms exposure) were recorded. When capturing images, we alternated between the two strains with ∼5 seconds between images, and the cells on one side of the chip were captured every 15 seconds. Each experiment included multiple runs, imaging 20-25 traps per side. To minimize artefacts, we started the experiments on different sides of the chip. We began image acquisition 10 seconds before removing IPTG (switching pressures to 240 mbar for M9 media and 90 mbar for IPTG), ensuring synchronized capture of equilibrium binding (following 2-minute IPTG exposure) and re-binding events post-IPTG washout. Experiments were performed at 30°C.

### Analysis of single-molecule experiments

#### Image analysis

Cells were segmented based on the phase contrast images using a U-Net convolutional neural network(37). Fluorescent dots (mVenus molecules) were detected using a wavelet-based algorithm(38): thresholding non-significant wavelet coefficients in the second á Trous wavelet plane. The threshold was defined as 5σ, where σ is the MAD estimate of the s.d. in the second wavelet plane(39). The average number of dots per cell versus acquisition time (beginning with an initial measurement taken under steady-state with IPTG present, followed by subsequent time points recorded after IPTG removal through media switch) was collected to obtain binding curves. The average number of dots detected in the strains that did not contain the *O_sym_* operator was first subtracted from the average number of dots detected in the strains with the *O_sym_*operator to avoid counting non-operator binding into specific operator binding. Because images from the two strains were recorded with a 5-second offset, interpolation was applied to align the data before subtraction.

#### Kinetic Analysis of LacI Binding to O_sym_ After IPTG Removal

To account for the sigmoidal shape of the binding curve, which is especially prominent in the Q55N LacI variant, we introduce a model in which IPTG efflux is explicitly modeled and IPTG binding to LacI inhibits it from switching into the specifically bound state. The model extends the three-state model from Marklund *et al.*(*27*), in which LacI can bind to the stretch of DNA that contains the *O_sym_* operator sequence. While on the DNA, LacI can slide and possibly bind specifically to the *O_sym_*, or dissociate from DNA. The extension to the Marklund *et al.* model lies in that LacI also has the possibility of binding IPTG. IPTG is assumed to be able to bind LacI while it is free and is non-specifically bound, but not when it is specifically bound to *O_sym_*. Limiting the number of DNA-bound states of LacI to only two, gives a manageable model with a manageable number of fitted parameters. We do, however, note that by excluding a state where LacI is on top of the Osym site, but not in its specifically bound conformation; a state to which IPTG can bind, will make the IPTG induced dissociation from the specifically bound LacI unrealistically slow in our model. We deemed this limitation of our model to have a small effect in our experiments where IPTG is excluded from the growth medium. In our model, IPTG binding is assumed not to change the properties of LacI non-specifically bound to DNA, and the non-specific binding to DNA is assumed not to change the IPTG affinity to LacI(40). IPTG binding and release are assumed to be rapid as compared to LacI-DNA binding, such that the IPTG binding reaction can be treated as at equilibrium. Taken together, the LacI binding can be described by the reaction scheme below, same as in Figure 4B:

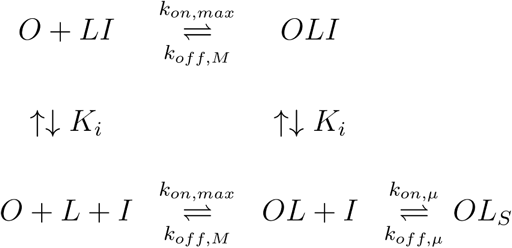

where:

● **O**: the specific operator site (*O_sym_*) in the genome.
● **L**: LacI protein
● **OL**: the complex of LacI bound to the operator non-specifically (corresponding to the testing state in the 3-states model stated in (27).
● **I**: IPTG
● **OL_S_**: the complex of LacI bound to the operator specifically
● *k_on,max_*: the on rate into testing state from unbound state to operator site
● *k_off,M_*: dissociation rate from the testing (non-specifically bound to the operator site) state to the state of unbound with operator site
● *K_i_*: IPTG binding constant
● *k_on,μ_*: the on rate of testing state LacI transition into the specific-bound state on the operator.
● *k_off,μ_*: the off rate of specific-bound state LacI go back into the testing state.

The decrease in the intracellular IPTG concentration is modelled using first-order kinetics, and, as in Marklund *et al.*, the non-specifically bound *OL*-state is assumed to be very sparsely populated, such that the specifically bound state and the IPTG concentration can be described by the following set of ODEs:

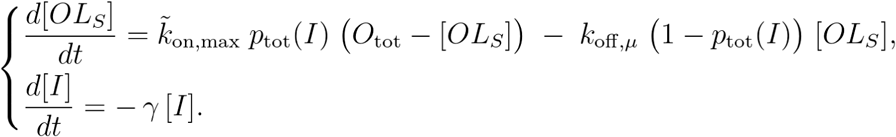

where 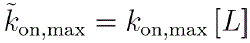 and the IPTG-concentration([I])-dependent probability of forming the *OL_S_* complex when in the *OL* state, O_tot_ is the total concentration of O_sym_ operator in the cell, 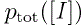 is

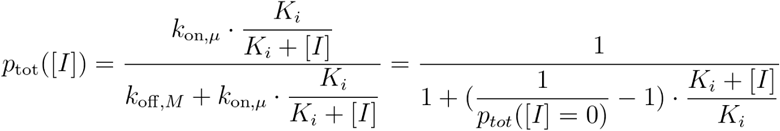

The free LacI concentration is assumed to be approximately constant throughout the experiment. In the regression, the following parameters are fitted: 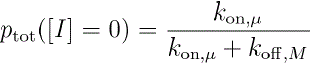, 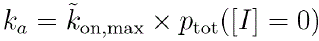, 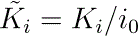, O_tot_, and γ. Here *i*_0_ is the initial IPTG concentration before the switch to media without IPTG. In the regression, the relation between association and dissociation rates is locked to the Miller measurements in the same strain using the relation

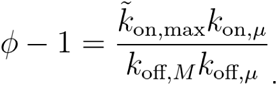

Note that the fitted in vivo association rate, k_a_, encompasses macromolecular crowding, 3D diffusion, 1D sliding/hopping, non-operator binding, and re-association; i.e., it reflects the net rate of forming a detectable specific complex at O_sym_ in vivo.

#### Miller assay

Overnight cultures were grown in M9-Glucose medium with and without 0.3 mM IPTG at 37°C. The overnight cultures were diluted (1:300) in fresh medium and grown to an OD_450_ of 0.5 – 0.6, where the exact value was recorded for each sample. For each combination of conditions (i.e +-IPTG) and LacI variant (*i.e.*, Wt, Q55N, and V52A), five replicates of 100 µL cell culture samples were prepared. Each culture sample was lysed in 700 µL Z buffer (60 mM Na_2_HPO_4_, 40 mM NaH_2_PO_4_, 10 mM KCl, 1 mM MgSO_4_, pH 7, with freshly added 10 mM DTT and 0.1% SDS) and 150 µL chloroform. The mixture was then vortexed for at least 5 seconds and incubated at 28°C for 5 minutes to ensure complete lysis. The enzymatic reaction of β-galactosidase was initiated by adding 100 µL of ONPG (4 mg/mL) substrate solution, allowing the yellow color (from the product o-nitrophenol) to develop. The reaction was terminated by the addition of 400 μL 1 M Na₂CO₃ at five different reaction stop times for each condition of a given LacI variant. In the presence of IPTG in the growth medium, the reaction times ranged from 5–63 minutes for the DelLacI strain, 13–72 minutes for the Wt-LacI strain, 10–69 minutes for the Q55N-LacI strain, and 21–143 minutes for the V52A-LacI strain. Without IPTG, the reaction times were longer, ranging from 6–65 minutes for DelLacI, 30–150 minutes for Wt-LacI, 12–71 minutes for Q55N-LacI, and 53–304 minutes for V52A-LacI. Shortly after each reaction was terminated, the mixtures were transferred to microfuge tubes, centrifuged at 12,000 x g for 1 minute, and the supernatants were collected for Abs₄₂₀ measurements.

Miller Units were calculated by determining the slope of Abs₄₂₀ values over reaction time using linear regression, normalized for culture density and sample volume. The formula used was: Miller Units = (1000 × SLOPE(Abs₄₂₀, Reaction Time)) / (Abs_450_ × 0.1), where the slope of Abs₄₂₀ values (reflecting β-galactosidase activity) versus reaction time was scaled by a factor of 1000 and divided by the absorbance at 450 nm (Abs_450_) to account for cell density, as well as the sample cell culture volume of 0.1 mL.

#### Representing Repression Strength and O_sym_ Occupancy via Miller Units

To isolate the repressive effect of LacI-venus variants, Miller Units for LacI-venus variants were normalized against the ΔLacI-venus strain (lacking LacI, hence completely non-repressed β-gal production). At steady state, the [β-gal] level depends on the operator’s occupancy: in LacI variants, [β-gal]=*γ*⋅*P_free_*/*μ* (where *γ* = maximal production rate, *μ* = degradation rate of β-gal), while in ΔLacI (where the operator is always free, *P_free_* = 1), [β-gal]=*γ*/*μ*. Normalizing Miller Units (which is proportional to [β-gal] in cells) gives: 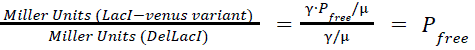, directly reflecting the probability of the lac operator (*O_sym_*) being unoccupied, *P_free_*= 1-*P_bound_* = normalized Miller Units. Then the inverse of normalized Miller Units (*ϕ*)*, ϕ* = 1/(1-*P_bound_*), shows the correlation with repression strength: higher *P_bound_* leads to higher *ϕ*. *O_sym_* occupancy can thus be quantified as 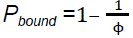.

### EMSA (Electricity mobility shift assay)

Electrophoretic mobility shift assays (EMSAs) were performed to analyze the difference in the affinities of *O_sym_* DNA with LacI-Halo variants (the Wt and Q55N mutant, same purified and protein labelled as in the PBM experiment stated above) in different salt concentrations. 1nM 5’IRdye700-labeled *O_sym_* containing 60bp DNA (same nt sequence as in PBM) were used in binding reactions with varying concentrations of LacI proteins (ranging from 0 nM to 500 nM) at different NaCl concentrations (20 mM, 100 mM, and 200 mM). Binding reactions were carried out in a 12 µL volume containing 1 nM DNA, 1x Imaging Buffer (same buffer as stated in the PBM experiment, except varying NaCl concentration). Negative controls included reactions without DNA or reactions without protein. The reaction mixtures were incubated at room temperature for 1 hour. Samples were loaded onto 5% Mini-PROTEAN® TBE gels (15 wells). The gels and running buffer(0.5X TBE) were pre-chilled and run at 200V for 35 mins to separate the protein-DNA complexes from free DNA. The gels were then imaged with Chemidoc (Bio-Rad) and analyzed with Imagelab for quantifying the LacI-bound fraction of DNA.

## RESULTS

### In silico Evaluation of Hinge Helix Propensity in LacI variants

In the LacI dimer the hinge regions connect the DNA-binding domains and the effector (allo-lactose/IPTG)-binding core domains. They form helices in the LacI crystal structure when bound to operator DNA (pdb id: 1efa) (44) but remain unstructured in the NMR structure, in which a truncated form of LacI with only the DNA binding domain and the hinge helices, bind to non-operator DNA(41). This structural plasticity in the hinge regions was hypothesised to drive the switching between the search and the recognition conformations (Fig. 1A)(45, 46). Guided by this hypothesis we selected five single amino acid substitutions that are expected (43) (Fig. 1C) to change the propensity of forming a helix and at the same time minimize unwanted side-effects in terms of protein structure of DNA binding. The helical propensities of an 18 amino acid peptide centered around the helical region of the hinge in the 1efa structure were analyzed using the Agadir software(47) (Figure 1D). As expected from intrinsic helical tendencies (Figure 1C), Q55N, Q54N and L56V have decreased helix propensity and G58A has increased helix propensity as compared to wt. The expected increase in helix propensity of Valine to Alanine mutation (Figure 1C) is primarily in the Arg51-Glu54 part of the helix. The hinge helix propensity of wt, Q55N, G58A, and V52A were in addition also estimated using all-atom molecular dynamics (MD) simulations. The Q54N and L56V were not selected for MD simulation due to the similarity to Q55N in the Agadir helicity estimates. Quantifying the transition between search and recognition states using all atom MD simulations of LacI bound to operator DNA is computationally intractable and we have previously shown (48) that enhanced sampling of the transition is non-trivial. Here we, instead, simulated the free LacI dimer in solution and quantified the number of hinges that form helices (defined as having an C-alpha RMS deviation from ideal helix of R51-A57 < 0.1nm) (Fig. 1E). The wt simulations were started from the 1efa (15) crystal structure with DNA and ONPF removed and the mutant simulations with the corresponding mutation introduced in the wt structure. Note that all simulations were performed five times, see methods for details. For all of the LacI variants, the fraction of hinges in helical form decreases over time for about 300ns until reaching a plateau. In line with previous MD simulations of V52A(48), the helical fraction of the hinge decreased slower for the V52A as compared to Wt. The helical fraction of the G58A mutant also decreased slower while the Q55N showed similar behaviour as wt. The reason for lack of effect in the Q55N mutant in the MD simulation as compared to the estimates for the peptide alone is currently unknown. We do not expect the lack of an effect in the helicity of the Q55N hinge to be due to force field parameter bias in the MD simulations, since previous results using a similar forcefield (49) have shown good agreement between estimates of helicity in estimates of helicity in MD and experiments experiments for short peptides. Rather the expected decrease for Q55N could be masked by the uncertainty estimate, based on the five replicates. Taken together these results indicate that Q55N has a bias towards the search conformation and the G58A and V52A are biased towards the recognition confirmation. Although G58A displayed increased helix propensity, the initial *in vitro* experimental screening revealed weak DNA binding ability of G58A (Fig. S9). These in vitro results are in line with observations by Suckow, J et al, 1996(50), who showed that substitutions at G58 abolish LacI function. Based on these results, the Q55N and V52A were selected as examples of mutants having decreased or increased hinge helix propensities and thus, expected either a deceased or an increased bias towards the recognition conformation of LacI. The G58A mutant was not selected for further investigations, although its limited binding raises the question of whether helix formation is the most critical parameter guiding binding.

### High-Throughput In Vitro Binding Kinetics Reveal Specificity–Stability Trade-Offs

To test how the selected mutations impact the LacI’s binding strength to different DNA sequences and hence its specificity, we first measured the *in vitro* association and dissociation rates of selected variants with 2,479 DNA sequences, including the natural *O_1_*, *O_2_*operators, the artificially strong *O_sym_* operator, and their single and double mutants using a protein-binding microarray (PBM)(27, 51) (Fig. 2). We mounted two PBMs in adjacent flow cells on a microarray slide (Fig. S7). Wt-LacI was flowed over one of the PBMs while a LacI mutant was flowed over the other. Both proteins were fluorescently labeled. The binding dynamics for each LacI protein were measured by introducing the reaction buffer with LacI molecules (association phase) and subsequently introducing the reaction buffer without LacI molecules (dissociation phase). This process allowed us to capture the time-dependent increase (Fig. 2A&B, association) and decrease (Fig. 2A&B, dissociation) in binding signals. Relative association rate (*k_a_*) were obtained by fitting association curves and normalizing the fitted parameter to Wt-LacI binding to *O_sym_* in the same experiment. Dissociation rate constants (*k_d_*) were extracted from separate fits of the dissociation curves (see details in methods).

The V52A-LacI mutant associates faster than the Wt-LacI to most mutated operator sequences (as shown in Fig. 2A&C and Fig. S5A). The deviation between V52A and Wt, on average, increases for sequences with lower association rates. The dissociation rate for V52A is slower than that for Wt from all sequences (Fig. 2D and Fig. S5B), with a similar fold-change between V52A and Wt for all sequences. As a consequence, V52A-LacI has lower specificity (Fig. 2F) and generally a lower equilibrium dissociation constant, *K_D_*, as compared to Wt-LacI (Fig. 2E and Fig. S5C). The shift towards higher affinity to the most of DNA sequences in the V52A-LacI mutant is consistent with the hypothesis that increased helix propensity gives more stable binding.

The Q55N-LacI mutant, where the hinge region is expected to be less helical as compared to wt, shows a slower association rate as compared to Wt-LacI for most sequences. Q55N-LacI shows faster dissociation from most operators compared to Wt-LacI, with a fold change similar for most sequences. The shifts in association and dissociation rate constants as compared to Wt for the Q55N mutant result in an increased specificity (Fig. 2F), but a reduced binding strength for all operators, including *O_sym_* (Fig. 2E). The slow association and rapid dissociation for the Q55N mutant cause many weak binding sequences to fall below our detection limit. The *in vitro* data reveal that Q55N-LacI has a higher affinity to one of the *O_sym_* single base-pair mutants than to the *O_sym_* operator itself (Fig. 2E). However, this *O_sym_* single base-pair mutant is absent from the wt *E. coli* genome; the closest sequence in the wt *E. coli* genome differs by 3 base-pairs from *O_sym_* (Table S4). In conclusion, if *O_sym_* were to be introduced into the wt *E. coli* genome, Q55N-LacI would still recognize it with the highest affinity.

The *in vitro* results agree with a conformation-switch model, in which the changes in preference for the search or recognition conformations impact the specificity and binding stability of LacI to *O_sym_*. Biasing the hinge region of LacI towards adopting the helical (recognition) conformation stabilises the binding on operator sequences, but it also elevates the binding probability on non-operator sequences, thereby reducing DNA specificity (and vice versa). The remaining question is how the propensity of forming helices in the hinge region of LacI affects its target DNA search speed in the intercellular context with all competing non-operator DNA sequences present in the genome. Based on extrapolation from the *in vitro* data, predicting the operator access time in relation to that of Wt-LacI for Q55N-LacI is straightforward. Q55N-LacI hardly interacts with any sequences more than 1 base pair mutation away from the *O*_s*ym*_. Since Q55N should have a lower probability adopting the recognition conformation on non-operator sequences the operator access time should be short, possibly shorter than that of Wt-LacI. However, the prediction for the total *O_sym_* occupancy *in vivo* is not as straightforward, because the *in vitro* dissociation rate of Q55N-LacI from the *O_sym_* operator is faster than that of Wt-LacI, which, together with potentially shorter operator access time, makes it hard to predict. The specificity of V52A-LacI is lower than that of Wt-LacI for near-operator sequences. This means that *in vivo,* the operator access time may be impacted by competing binding to non-operator sequences, *i.e,* the operator access time could be longer than that for Wt-LacI. However, dissociation from *O_sym_* is slower, again making the *in vivo* steady-state binding uncertain without live-cell measurements.

### In Vivo Measurements Reveal Weak Coupling Between Specificity and Search Speed

To measure the *in vivo* steady-state binding of the LacI variants to *O_sym_*, we constructed strains in which the expression of β-galactosidase (β-gal) is regulated by LacI binding to *O_sym_*(Fig. 3A). The intracellular β-gal concentration can then be measured using the Miller assay and thus be used to estimate the steady-state LacI binding to *O_sym_*. We determined the *in vivo* steady-state binding to *O_sym_* with the Miller assay(52) for β-gal activity using the same halo-tagged dimer version of LacI as in the *in vitro* experiment. If the repressor is bound at the operator, the expression is off, which makes the activity proportional to the fraction of time the operator is unoccupied(53).

**Figure 3.**
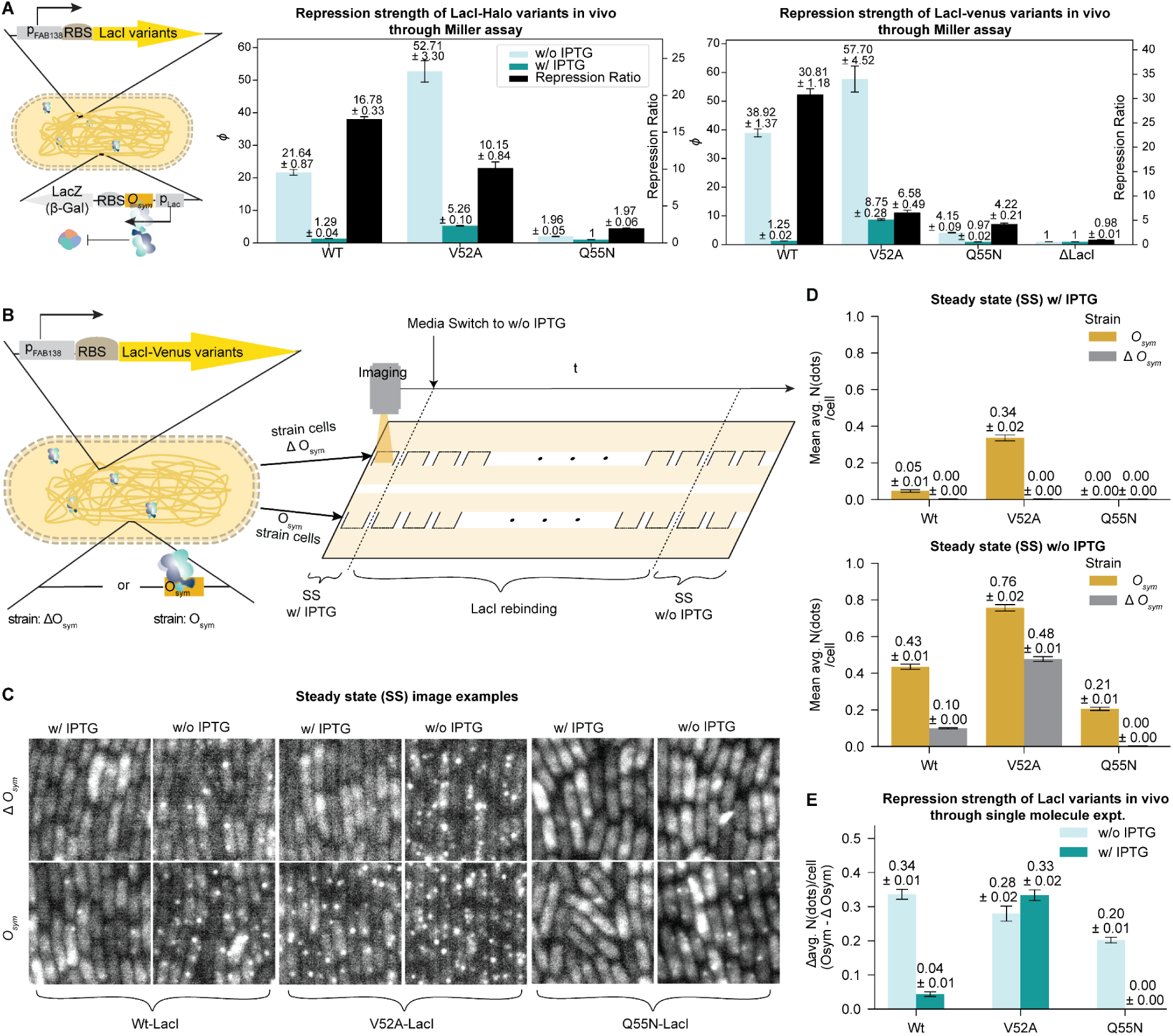
Steady state binding of LacI variants *in vivo* (A: Miller assay, B-E: Single-molecule experiments). **A**. Repression strength (1/ Normalized Miller Units, denoted as *ϕ*) for LacI-halo variants (*middle*) and LacI-venus variants (*right*) through Miller assay. The β-gal activity (Miller Units) was measured under the control of the *O_sym_* operator with three LacI protein variants expressed from the *E. coli* genome (*left*). *ϕ* indicates the repression strength of each LacI variant on *O_sym_*. For the LacI-halo strains, as shown in the left bar plot, the Q55N variant with IPTG was used as a control, representing a state of no repression (where *ϕ* = 1). For the LacI-venus strains, as shown in the right bar plot, the ΔLacI variant served as the control for each state without (w/o) IPTG or With (w/) IPTG. All LacI variants were tested in 3 biological replicates. Data are presented as Mean ± SEM **B.** A schematic overview of the single-molecule experimental workflow conducted on a microfluidic chip. Two isogenic *E. coli* strains, both expressing a certain Venus-tagged LacI variant but differing in their genomic presence or absence of the *O_sym_*operator sequence, were independently introduced into hydraulically interconnected cell chambers (labelled as squares open on one side), which are positioned on opposite sides of the chip. This setup allows strains to have simultaneous access to the shared growth media. Initially, chambers were exposed to media containing 0.3 mM IPTG. Fluorescence imaging was performed to capture the steady-state (SS) LacI binding in each strain under the IPTG-supplemented condition (SS w/ IPTG). Following this, media was exchanged to IPTG-free media, and time-lapse fluorescence imaging subsequently captured downstream chambers on both sides of the chip to quantify LacI rebinding kinetics until a new steady-state equilibrium was established in media without IPTG (SS w/o IPTG). **C.** Fluorescence images of cells expressing three LacI variants at steady state under two conditions: without IPTG (w/o IPTG) and with IPTG (w/ IPTG). Columns correspond to the IPTG condition, and rows correspond to two genetic backgrounds: one with the *O_sym_* operator sequence integrated into the genome and one in which this operator is not present (Δ*O_sym_*). For each LacI variant, the paired images of the *O_sym_* and Δ*O_sym_* strains are indicated by brackets. **D**. Detected average number of binding dots per cell for LacI variants in strains with or without *O_sym_*, in the steady states with IPTG (*Top*) or without (*Bottom*). **E.** The *in vivo* repression strength of LacI variants to the *O_sym_* operator is expressed as the average number of specifically bound LacI molecules, which is calculated by subtracting the average number of dots per cell in the Δ*O_sym_* control strain from that in the *O_sym_* containing strain measured at steady state, both with IPTG (w/ IPTG) and without IPTG (w/o IPTG).

The Miller units are directly proportional to the intracellular concentration of β-gal. For the strains carrying the *O_sym_* operator in the genome, the relationship between measured Miller units and the chance of the *O_sym_* operator being bound by LacI can be written as: Miller units ∝ [β-gal] ∝ *P_free_* (probability of *O_sym_* being free). *P_free_* can be calculated as 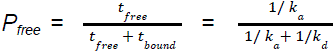 and the inverse of the normalized Miller units, abbreviated as *ϕ*, is then equal to 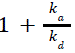. This implies that the ratios of *ϕ* correlate to the ratio of affinities (1/*K_D_*), such that 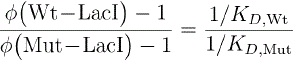, assuming similar free intracellular LacI concentrations across the variants (controlled by the same promoter).

To estimate the *in vivo* association rate based on the steady-state binding estimate from the Miller assay, we need an estimate of the dissociation rate. If the relative dissociation rates of Wt-LacI and V52A-LacI observed *in vitro* (Fig. 2E) are similar to *in vivo*, the dissociation rate of V52A-LacI would be roughly 2.4 times slower than that of Wt-LacI. *In vivo*, V52A-LacI repressed the expression of β-gal approximately 2.5, (52.71-1)/(21.64-1), times stronger than Wt-LacI (Fig. 3A left bar plot). The close agreement between these values suggests that V52A-LacI may search equally fast as Wt-LacI *in vivo*. This would imply that V52A-LacI is not slowed down much by interactions with non-operator DNA or that it binds with a higher probability when it reaches the *O_sym_*operator *in vivo*. Importantly, but independently of the search speed, V52A-LacI represses β-gal expression to a large extent also in the presence of IPTG, where the expression of β-gal is only 11% of that of the ΔLacI strain.

For the Q55N-LacI mutant, the binding strength to *O_sym_* compared with Wt-LacI *in vivo* decreases by a factor of ∼20, while its *in vitro* dissociation rate only increases by a factor of 3. Assuming that the relative dissociation rates of Wt-LacI and Q55N-LacI remain similar *in vivo*, and that the Q55N-LacI access time to *O_sym_* is shorter or similar to Wt-LacI, due to a lower probability to take the recognition conformation on the non-operator sequences, then the search time of Q55N must be extended as compared to Wt by a lower binding probability to *O_sym_* than what is measured *in vitro*.

To directly visualize the *in vivo* steady-state binding results, as assessed in the Miller assay above, and also to measure the time it takes for the LacI variants to access and specifically bind to the *O_sym_*operator *in vivo,* we used the single-molecule method we previously described in Hammar *et al.*(*6*). Following Hammar *et al.*, we expressed LacI as a translational fusion to the fluorescent protein Venus at only a few molecules per cell in a strain carrying a single *O_sym_* in the genome (Fig. 3B). The operator is not regulating any promoter in this strain. The exact number of LacI-venus molecules in the cell is not critical as long as it is the same for the different variants, and the total fluorescence from freely diffusing molecules is sufficiently low to allow the observation of one chromosomally bound dimer as a diffraction-limited fluorescent spot when we use a long (4s) exposure time. The rationale is that only the specifically bound molecules give a signal over the background since the emission light from the non-bound molecules is blurred by diffusion. The bacteria are grown in a microfluidic device that allows for rapid media switches while imaging the two types of growing cells on the microscope (Fig. 3B)(54).

First, we quantified the steady-state (SS) *in vivo* binding of each LacI variant under two IPTG conditions—with IPTG (SS w/ IPTG) and long after IPTG removal (SS w/o IPTG)—in two genomic backgrounds: one with the *O_sym_* operator (strain type: *O_sym_*) in the genome and one without (strain type: Δ*O_sym_*), as illustrated in Fig. 3B. Example images taken in steady-state are shown in Fig. 3C for different LacI variants. When spots are observed in strains without the specific operator sites, these correspond to long-lasting (>4s) binding to non-operator sequences. Fig. 3D shows the corresponding quantification of the average number of detected binding spots per cell, under steady-state conditions with or without IPTG. In SS w/ IPTG, Wt-LacI shows no spots in the Δ*O_sym_* strain and a small number of spots due to specifically bound LacI in the *O_sym_* strain. In SS w/o IPTG, there are some binding on non-*O_sym_* DNA sequences in the Δ*O_sym_*strain, but the *O_sym_*strain displays more binding events. V52A-LacI in SS w/o IPTG displays more binding to non-*O_sym_* DNA sequences than the other two LacI variants. This is in line with the stronger binding of the V52A-LacI to operator mutants *in vitro* as compared to Wt-LacI. While IPTG reduces the V52A-LacI binding below the detection limit when there is no *O_sym_* in the genome, it is less effective at reducing the number of bound LacI molecules when *O_sym_* is present in the genome (Fig. 3D *left*). The large *O_sym_*occupancy by V52A-LacI even in the presence of IPTG corroborates the Miller assay results, with which the *O_sym_* occupancy can be calculated as 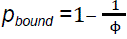, see derivation in methods. The calculation gives a probability of 89% (*i.e.* 1 – 1/8.75 ≈ 89%) that *O_sym_*is bound by V52A-LacI in the presence of IPTG, this loss of inducibility phenotype for V52A mutant was also reported in previous studies(50, 55). Q55N-LacI, on the other hand, only displays binding events when there is *O_sym_* in the genome and is fully dissociated by IPTG. This is also expected from the *in vitro* experiments and the Miller assay data.

Next, we evaluated the dynamics of target search for Wt-LacI and Q55N-LacI just after removing IPTG (through media switch) in single-molecule experiments. The experiment could not be performed for V52A-LacI, because the *O_sym_*occupancy by V52A-LacI remained high even with IPTG present, as seen in Figure 3E. Any binding increase after IPTG removal likely comes from its binding to non-operator sequences. The Miller assay also shows that *O_sym_*occupancy in the V52A-LacI strain increases from 89% with IPTG to 98% (1-1/*ϕ* = 1 – 1/57.7 ∼ 98%) without IPTG. In contrast, for the *O_sym_*operator, occupancy by Q55N-LacI is changing from close to zero to 59% of that of Wt-LacI, as shown in Figure 3E (0.20/0.34 ≈ 59%). Therefore, by removing IPTG from the growth medium, we can directly measure the Q55N-LacI binding rate using time-lapse imaging, similar to our previous studies with Wt-LacI(3, 6).

In Figure 4A, binding of Wt-LacI and Q55N-LacI to the *O_sym_*operator over time after IPTG removal is shown as the difference in the average number of detected spots per cell in strains with *O_sym_* in the genome compared to the Δ*O_sym_* strain. To analyze the specific binding kinetics of the LacI variants to the genomic *O_sym_*site after IPTG removal, we used a version of the kinetic model used in Marklund *et al.*(*27*), now extended to include IPTG. In this extended model(Figure 4B), LacI is not allowed to form the specifically bound state if it is bound to IPTG (see more details in methods). The change of intracellular IPTG concentration in the model continuously decreases due to the net IPTG efflux after switching to media lacking IPTG. LacI is controlled under the same promoter as in the Miller assay, so similarly, a constant concentration of free LacI throughout the experiment is assumed. Binding affinities measured using the Miller assay are assumed to be valid also for the *in vivo* kinetics measurement, because the ratio of *O_sym_* occupancy between Q55N-LacI and Wt-LacI is measured to be in close agreement in the Miller assay and the single-molecule experiments. In the Miller assay, the *O_sym_* occupancy for Q55N-LacI to Wt-LacI is (1–1/4.15)/(1–1/38.92) ≈ 0.78. In the single-molecule experiments, the ratio is calculated as above, 0.20/0.34 ≈ 0.59. With the two assumptions above, we fitted the Figure 4B model to the binding data in Figure 4A, allowing us to estimate the average *k_a_* to the *O_sym_* operator *in vivo*. We find that the average time for a single *O_sym_* operator in the bacterial genome to be found and bound by LacI is around 0.7 minutes in both the Wt-LacI and the Q55N-LacI containing strains (Figure 4D). This indicates that making LacI prefer the search conformation does not accelerate its association rate to the operator site by minimizing non-operator interactions, as we initially anticipated. Instead, the primary effect of making LacI prefer the search conformation appears to impact much more on the dissociation rate with its target operator: since Q55N-LacI exhibits a 20-fold weaker *in vivo* binding to *O_sym_* compared to Wt-LacI (as shown by the Miller assay), yet maintains a similar association rate in single-molecule experiments *in vivo*, it follows that the difference in binding strength arises from changes in the dissociation rate. However, the ∼3-fold difference in *O_sym_*dissociation rates between Q55N-LacI and Wt-LacI observed *in vitro* is insufficient to fully explain the 20-fold change in *in vivo O_sym_*binding affinity, a discrepancy that is further discussed in the Discussion section.

**Figure 4.**
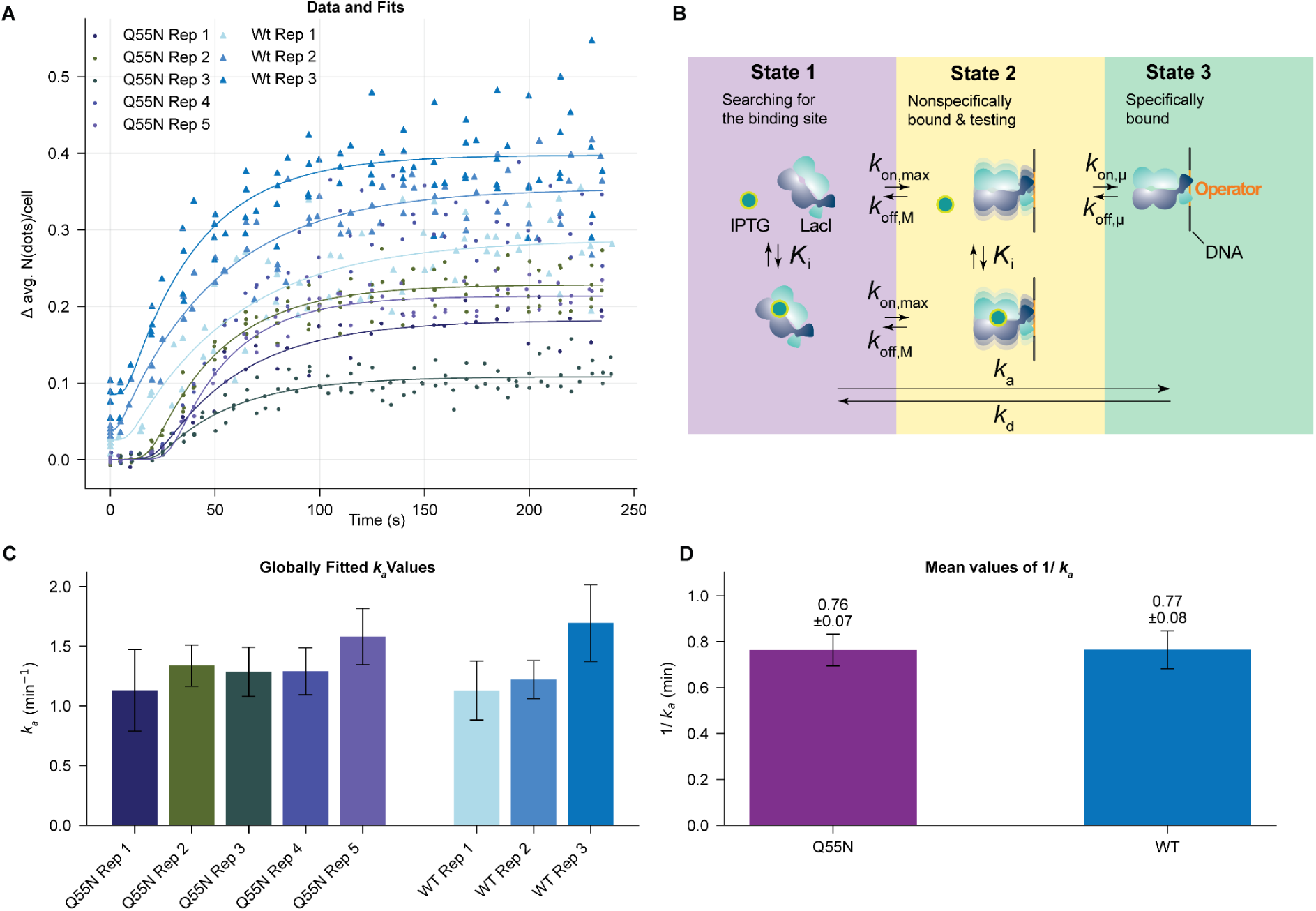
Binding kinetics of LacI variants to the *O_sym_* operator *in vivo* over time after IPTG removal. **A**. Binding is quantified by the difference in fluorescent spots per cell between *O_sym_*-containing strains (*O_sym_*) and control strains lacking *O_sym_* (Δ*O_sym_*), in the same way as in Fig. 3E. The binding curves were fitted using the IPTG-integrated three-state model (solid lines; see model in panel B and details about fitting in Methods). **B**. Three-state model, taken from Marklund *et al.*(*27*), integrated with IPTG. Microscopic rates entering and exiting from one state to another are labelled at the cross sections of states, *K_i_* is the equilibrium dissociation constant for the interaction between IPTG and LacI that is either in state 1 or state 2. Macroscopic association rate (*k_a_*) and dissociation rate (*k_d_*) are labelled out as for a LacI molecule from being far away from its target operator to being specifically bound to its operator. Parameterization and fitting are described in Methods. **C**. Bar plots of fitted parameter: *kₐ* value with SE for each individual replicate for Wt-LacI and Q55N-LacI, respectively. **D**. Average *in vivo* search times for Q55N-LacI and Wt-LacI molecules in a cell targeting a single *O_sym_* operator in the bacterial genome. This time is calculated as the inverse of the averaged fitted *k_a_*, with error bars showing the propagated standard error.

## DISCUSSION

In this study, we investigated whether the relative stability of the search and recognition conformations of the transcription factor LacI gives the predicted impact on search speed and binding strength, under the assumption that the evolutionary purpose of the conformation switching is to resolve the speed-stability paradox by having one conformation rapidly searching and the other conformation stably binding. Previous work identified the hinge region of LacI as critical for transitioning between the two conformations (42). We hypothesized that mutations in the hinge region would alter the search or recognition conformation preference, thereby impacting DNA binding stability and specificity, with the associated impact on target search kinetics in the genome. Using MD simulations, we identified a LacI mutant: V52A-LacI, which is shown in MD to favor a hinge region in the helix conformation and thus presumably favour the recognition conformation. In addition, using empirical predictions for short peptides (47), we identified that Q55N-LacI favor the unstructured conformation, thus presumably favouring the search conformation. *In vitro* DNA microarray experiments confirmed these predictions, with V52A-LacI showing stronger, but less specific, binding and Q55N-LacI exhibiting higher specificity, but weaker binding.

Experiments on living cells further demonstrated that V52A-LacI repressed a reporter protein controlled by *O_sym_* stronger than Wt-LacI (Figure 3A). The change in repression strength quantitatively corresponded to the slower dissociation rate for V52A-LacI *in vitro* (Figure 2G), suggesting the search time for the *O_sym_* site *in vivo* should be similar for V52A-LacI and Wt-LacI. At the same time, we observe a 4.8-fold increase in binding to non-operator sequences in the absence of a specific operator for V52A-LacI as compared to Wt-LacI (Fig 3D). Why do these non-specific binding events not substantially increase the search time for the V52A-LacI mutant? If we assume that the dissociation is equally impacted for Wt and V52A-LacI going from *in vitro* to *in vivo*, the most straightforward explanation is that V52A-LacI has a significantly higher probability of binding when reaching the *O_sym_* operator than Wt-LacI. However, this explanation seems unlikely considering that we found the probability of binding for V52A to be very similar to that of Wt-LacI *in vitro*, as shown in Figure S1B. Other potential factors underlying this discrepancy are explored below.

For the Q55N-LacI mutant, the time for searching the *O_sym_*site *in vivo* is similar to Wt. Following the same argument as for the V52A-LacI mutant, the time to access *O_sym_* could actually be shorter if the probability of binding is decreased. In the *in vitro* experiment (Figure S1B) we find that the probability for Q55N-LacI is decreased by 30%. If this is directly applicable in the *in vivo* situation this would imply a 30% decrease in the time to access to keep the search time the same as Wt. Q55N-LacI exhibited ∼20-fold weaker binding than Wt-LacI in the *in vivo* Miller assay (Figure 3A), which is only partly explained by the 3-fold faster dissociation observed *in vitro* (Figure 2G, see Table S5 for an integrated overview of comparative parameters across *in vitro* and *in vivo* experiments). The time for searching, which we can measure directly *in vivo* (Figure 4), does not account for the remaining factor needed to get the low repression observed in the Miller assay. Clearly, some aspect of the *in vivo* situation is not accounted for. What could these missing aspects be?

There is a subtle difference between the strains used for the Miller assay (Figure 3) and those used for the *in vivo* kinetics (Figure 4). The strains used for the Miller assay, by necessity of the assay, have a promoter in front of the *O_sym_* operator, while the strains used for *in vivo* kinetics do not. A reduced binding probability *in vivo*, as measured by the Miller assay, can, for example, be due to competition with RNAP (RNA polymerase) or active displacement of Q55N-LacI by RNAP. However, this possibility is ruled out by the close agreement between the ratio of *O_sym_* occupancy between Q55N-LacI and Wt-LacI, obtained from the Miller assay (0.78) and single molecule experiments (0.59). This suggests that the presence of a promoter in front of *O_sym_* does not dramatically decrease the overall *O_sym_* occupancy by Q55N-LacI compared to Wt-LacI in the Miller assay.

Another potential missing aspect could be differences in the ionic strengths used in the *in vitro* experiments and the *in vivo* situation inside living cells, where the sum of Na^+^ and K^+^ is 30 mM *in vitro* and 200-300 mM *in vivo*. However, EMSA equilibrium binding comparisons reveal that the Q55N-LacI affinity is less sensitive to changes in salt concentration as compared to Wt-LacI. Specifically, while both proteins show an increase in *K_D_* from 50 mM to 200 mM NaCl, the increase is substantially greater for Wt-LacI (∼10-fold) than for Q55N-LacI (∼2-fold) (Fig. S8 C&D). Based on the EMSA result we would expect the Q55N-LacI and Wt-LacI to become more similar going from *in vitro* to *in vivo* and not vice versa which is the observed. The lower sensitivity of the Q55N-LacI affinity to salt increase as compared to Wt-LacI is qualitatively inline with a model where increasing salt concentrations primarily decreases the sliding length (as detailed in the SI).

Macromolecular crowding is present in the *in vivo* not in the *in vitro* experiments. Crowding could affect the *in vivo* situations in multiple ways: For bi-molecular interactions in 3D, obvious effects of crowding are increased concentration due to excluded volume and changes in diffusion rates. These effects will, however, most likely impact the three variants equally. A more subtle effect of crowding is that it promotes compacted protein conformations in order to increase the entropy in the solvent (56), which may impact the LacI variants differently. Crowding would, to first approximation, promote the more compacted helix conformation. This would presumably make Q55N-LacI more similar to Wt *in vivo* as compared to *in vitro*, *i.e.* it cannot explain why Q55N-LacI binds worse *in vivo*. When there is crowding on DNA (i.e 1D) the free sliding length is restricted by other DNA-binding proteins (57). These reduced sliding lengths make the search slower, especially for proteins that slide far. On the other hand, the crowding also causes more return events to the operator after dissociation from the operator into the sliding mode. This leaves the equilibrium binding essentially unaltered and thus doesn’t explain the difference between *in vitro* and *in vivo* affinities. Lastly, the crowding on DNA also decreases the amount of accessible non-specific DNA and the fraction of time the operator is available, but these consequences are very similar for all variants. In summary, crowding cannot account for the differences between *in vitro* and *in vivo*.

Finally, a possible explanation could be that the chromosome supercoiling or dynamics make dissociation faster *in vivo* than on the short oligos presented in the *in vitro* experiment. If this effect is more pronounced for Q55N-LacI, which has a lower binding energy compared to Wt-LacI, then for V52A-LacI, which has a higher binding energy as compared to Wt-LacI, this could explain both why Q55N-LacI binds *O_sym_* weaker than what is expected from its *in vitro* dissociation rate and why V52A-LacI binds as expected. However, it is still not clear why V52A-LacI did not spend more time in the search while it clearly showed more binding than other LacI variants to non-operator sequences *in vivo*.

Independent of what causes the discrepancy between the *in vitro* and *in vivo* observations, we find that Q55N-LacI represses poorly *in vivo*, although it associates with the operator as fast as Wt-LacI. This observation has important implications regarding our initial question, whether favoring the search conformation can make the association faster. It appears the answer is no; the interactions between Wt-LacI and non-operator DNA sequences are already sufficiently low not to get trapped on non-operator sequences for significant fractions of time.

If we instead tune LacI to favor the recognition conformation, we find that V52A-LacI enhances repression strength mainly by making dissociation slower, but at the same time, it loses inducibility. Taking into consideration that the V52A mutant has shown an IPTG affinity comparable to Wt-LacI(55), the likely explanation for the reduced inducibility is that V52A-LacI binds so stably to the *O_sym_* operator that it only rarely flips out of the specific bound conformation, limiting the opportunity for IPTG to bind V52A-LacI. Increased binding strength to the operator together with a reduced inducibility as compared to Wt-LacI has also previously been observed for the V52 mutants, V52A, V52H and V52S when binding the *O_1_* operator. However, Zhan *et al.*, 2006 reported that tetrameric V52 mutants did not enhance affinity for *O_sym_*, whereas our measurements with dimeric LacI variants measurements show a clear affinity increase for *O_sym_* both *in vitro* and *in vivo*. The divergence, likely rooted in oligomeric state or assay design, does not alter the qualitative message shared by both studies: hinge-helix stabilization tightens DNA binding at the cost of IPTG responsiveness. We therefore attribute diminished inducibility to a shift of the conformational ensemble toward the recognition state, limiting access to the IPTG-accessible search state. This stability-inducibility trade-off is in line with earlier reports for mutations in other LacI domains(14, 58).

So how do we reconcile that G58A-LacI displayed increased helix propensity as compared to Wt-LacI, based on theoretical predictions, and that it at the same time binds poorly (50), Fig. S9? If switching between sliding and recognition confirmations is critical for facilitated diffusion, a transcription factor that is locked to the recognition confirmation should bind very slowly. For example, the recognition conformation may require a bent DNA structure, thus limiting binding to or sliding on straight DNA. G58A may represent such a locked recognition state. What speaks against this idea is that G58A still binds at a reasonable rate (Fig S9).

Circling back to our initial question, this supports that the operator access time can be traded for bound complex stability, but that the association rate (search speed) is not increased by faster access since it also impacts the binding probability at the operator. Our observations instead support the model in which the mechanistic challenge is to compromise between inducibility and repression strength. While it remains unclear whether Wt-LacI in its native context represents an evolutionary optimum with respect to this compromise, the existence of previously reported mutants(58, 59) with both enhanced repression and IPTG inducibility suggest that it is not. IPTG is, however, not LacI’s natural inducer and, therefore, it will be important to investigate whether these mutants keep enhanced repression and inducibility when interacting with the natural inducer (allolactose) in living cells to know if we still are missing pieces for understanding the evolutionary constraints shaping LacI function.

## ACKNOWLEDGEMENT

We are grateful to Jimmy Larsson for assistance in microfluidics experiments, to Anna Knöppel for assistance in strain construction, to Anneli Borg for the discussions on troubleshooting the PBM experiments and to Irmeli Barkefors for critical reading of the manuscript.

## AUTHOR INFORMATION

### Author notes

These authors contributed equally: Jinwen Yuan, Malin Lüking

### Authors and Affiliations

**Department of Cell and Molecular Biology, Science for Life Laboratory, Uppsala University, Uppsala, Sweden**

Jinwen Yuan, Malin Lüking, Spartak Zikrin, Beer Chakra Sen, David Fange, Johan Elf

**Department of Biochemistry and Biophysics, Science for Life Laboratory, Stockholm University, Stockholm, Sweden**

Emil Marklund

### Contributions

J.E. conceived the project; J.E., D.F. and E.M. supervised the project; M.L. and D.vdS. performed MD simulations; J.Y. designed the PBM study, purified and labelled LacI and carried out PBM experiments; D.F., J.Y., and S.Z. developed the theoretical models and data-analysis methods for PBM. J.Y. analyzed the PBM data. B.S., M.L., and J.Y. generated the strains used for both the Miller assay and SM experiments; M.L. and J.Y. conducted the SM experiments and, with S.Z.’s assistance, analyzed the resulting data; J.Y., B.S., and M.L. performed the Miller assays, and J.Y. handled the data analysis. D.F. conceived the kinetic model to fit SM binding data alongside the Miller assay results.; J.E., J.Y., M.L and D.F. wrote the paper, with input from all authors.

### Corresponding authors

Correspondence to Johan Elf

