## Supplemental Information for "The LacI hinge region balances binding stability against inducibility"

### Supplementary Text

*Derive Microscopic binding rates from measured macroscopic rates ($k_a$, $k_d$) on PBM*

To dissect the binding properties of the mutants at a more detailed level we adapted the theoretical approach from our previous study(Marklund et al. 2022). The approach uses the macroscopic measurements $k_a$ and $k_d$ to many operators to make inferences about microscopic properties in special cases. In particular, if there is a linear anticorrelation between $k_a$ and $k_d$ then the operator binding is mainly governed by the probability, $p_{\text{tot}}$, of specifically binding when the TF is non-specifically bound on the operator, and less by the rate of switching back to search confirmation, $k_{off,u}$, when bound on the operator. For the V52A mutant, the anticorrelation trend is clear (Fig S1A), which makes it possible to infer its microscopic binding rates (see Material and Methods for details). Importantly, the microscopic rate of V52A for leaving the bound state is lower than Wt, which is consistent with that it prefers the helix conformation of the recognition complex. The probabilities for binding operators, $p_{tot}$, (Fig S1B) are also generally higher than for Wt For Q55N-LacI the specificity is so high that there is little data to apply the model to. This implies that $p_{\text{tot}}$ are very low, except for $O_{sym}$ which is still close to unity. For the operators where the binding and unbinding events are still possible to measure, there is an anticorrelation for $k_a$ and $k_d$ that suggests that the microscopic rate of switching back to the search confirmation is slightly higher than Wt-LacI (Fig S1D).

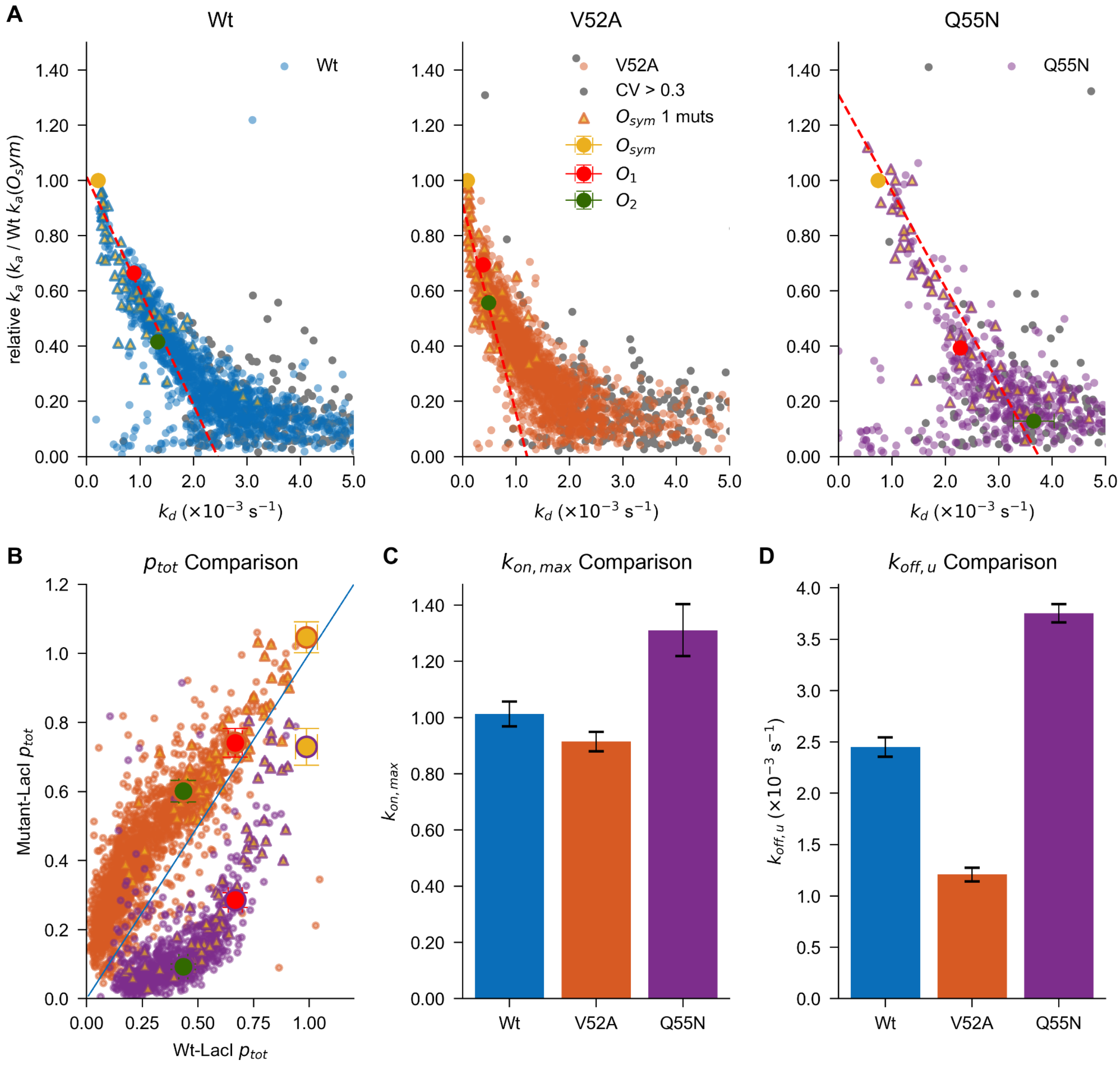


**Figure S1. Measured $k_a$ vs $k_d$ rates for LacI variants binding with thousands DNA operators *in vitro*.** **A.** Measured association and dissociation rates for LacI-Wt (Blue), LacI-V52A (Orange) and LacI-Q55N (Purple) binding with thousands of single and double mutants of three DNA operators of LacI ($O_{sym}$, $O_1$, $O_2$) *in vitro*. Each data point represents the mean $k_a$ value for a unique DNA sequence, calculated from 10 replicates for Wt-LacI and 5 replicates for each mutant-LacI. Red dashed lines show the model prediction of $k_a = k_{on,max} - k_d * k_{on,max}/k_{off,\mu}$ , where $k_{on,max}$ is the intercept of a linear fit to the top 35% of $O_{sym}$ single mutants (CV < 0.3) and $k_{off,\mu}$ is the trimmed-mean microscopic unbinding rate estimated from all $O_{sym}$single mutants (see details in methods). Three LacI operators ($O_{sym}$, $O_1$, $O_2$) are labeled with yellow-, red-, and green-filled circles, as marked. Excluded sequences associated with weak binding or high variability (CV ≥ 0.3) of $k_a$, $k_d$ measurement are marked in grey points as in Figure 2. Error bars on three operators are propagated standard error of mean (S.E.M.) of their fitted and normalized $k_a$ and fitted $k_d$ measurements. **B.** Calculated binding probability for Wt-LacI(X-axis) vs. mutants (Y-axis), according to Marklund et al.(Marklund et al. 2022) for each protein ($k_a$ / $k_{on,max}$), with y=x as blue line, representing Wt-LacI's $p_{tot}$ for comparison. **C & D** The $k_{on,\,max}$ value is determined as the y-intercept from a linear fit performed on the top 35% of Osym single mutants, with its error bar representing the standard error (SE). The mean and SEM for $k_{off,u}$ are calculated from a set of trimmed $k_{off,u}$ values—excluding the 10% most extreme measurements—from all obtained koff,u data. These $k_{off,u}$ values are computed by applying the equation $ka = kon,max - \frac{kon,max}{koff,u} \times kd$ (as presented in Eq. 1 of Marklund et al.(Marklund et al. 2022)) to all valid Osym single mutants (depicted as triangles with yellow face colors in panel A) that have an estimated $k_{on,max}$ from the linear fit.

#### Salt dependence of $K_D$

How can we reconcile that the $K_D$ of Q55N-LacI is less sensitive to ionic strength when compared to Wt-LacI? If we break down $K_D$ into components, we can disentangle which factors may respond differently for mutant and Wt LacI.

$$K_D = \frac{k_{off,\mu}(1 - p_{tot})}{k_{on,max} p_{tot}}$$ (Marklund et al. 2022 Eq S18)

$k_{off,u}$ is different for the different mutants (Fig. S1), but since it is mainly due to factors inside the protein DNA complex, it is unlikely to change differentially with respect to salt concentration. $k_{on,max}$, the rate of a LacI protein entering into the testing state around the DNA region that contains $O_{sym}$, can potentially change differently for the different proteins, but it's not clear how. The impact of a change in ionic strength on $p_{tot}$, the probability of binding the operator before leaving that region of the DNA, is, however, easier to predict. To simplify the notation, we defined $(1\text{-}p_{tot})/p_{tot}$ ($=q$), *i.e.,* a low $q$ ($=1/p_{tot}-1$), means a low $K_D$, which means a strong binding.

In a model where LacI slides in 1D on non-specific DNA before it macroscopically dissociates to an uncorrelated part of the chromosome, it returns to the same position on the DNA as many times as the number of base pairs in the sliding length(Berg et al. 2016). Thus, if we assume that the sliding length is N, $p_{tot}$ is one minus the probability of failing to bind N consecutive attempts, *i.e* $p_{tot}=1-(1-p)^N$, where $p$ is the probability of binding when the LacI is on top of the operator.

Based on previous studies, partly on other proteins, the ionic strength has a major effect on non-specific binding(Wang et al. 1977) and sliding length(Blainey et al. 2006) but not diffusion rate(Blainey et al. 2006). We can therefore evaluate what would be the effect on the binding constant, $K_D$, if we change the sliding length for proteins with different probabilities of binding when on top of the operator, $p$, assuming that $p$ is not dependent on salt concentration. In Figure S2 we plot the ratio of $K_D$, $Q=(1-p)^N/(1-(1-p)^N)\ /\ ((1-p)^n/(1-(1-p)^n))$, where N is the low salt sliding length (≈60) and n is the high salt sliding length (≈30).

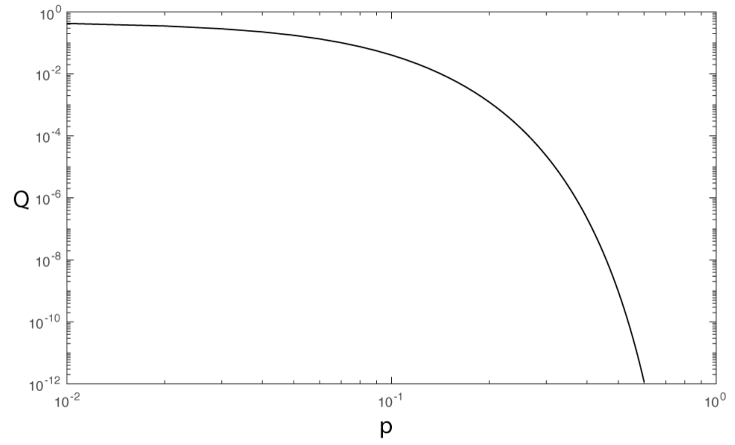


**Figure S2**

If we estimate $p_{tot}$ from Fig S1b we get $p_{tot}$= 0.99 for WT and $p_{tot}$ = 0.8 for Q55N, which translates to $p$=0.074 for WT and $p$=0.027 for Q55N, if we assume $N$=60 corresponding to the longer sliding in low salt. In terms of change to higher salt with shorter sliding, $n$=30, this gives $Q$=0.09 for WT and Q=0.31 for Q55N.

### Supplementary Figures

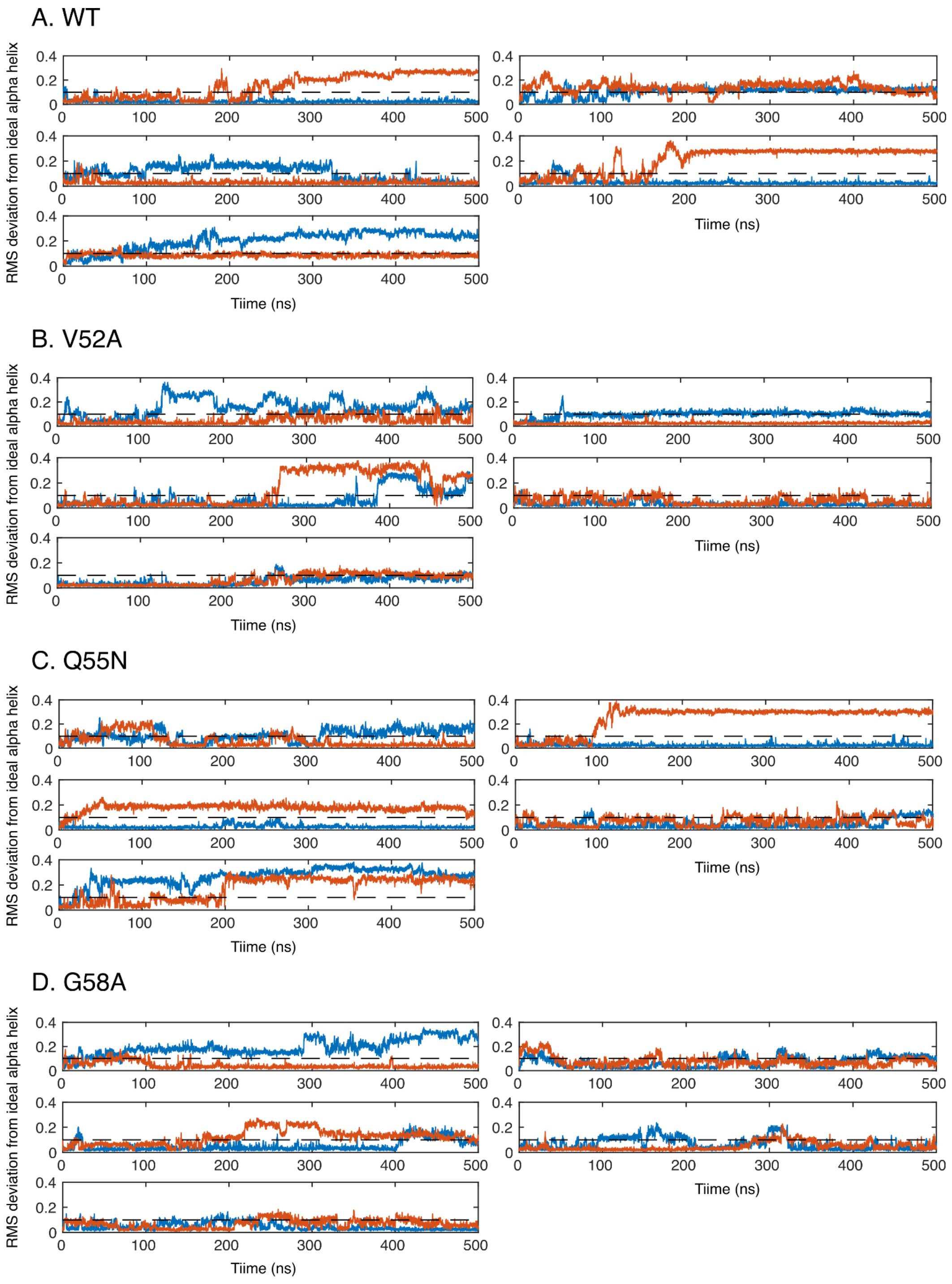


**Figure S3: Root mean square deviation from ideal helix for residues R51-A57 as function of time for five replicate MD simulations of a LacI dimer.** Each panel for each LacI variant contains results from one MD simulation. Red and blue lines show RMSDs for each of the two hinges in the LacI dimer. Black dashed line is RMSD=0.1nm, which is used as cut-off for assigning a hinge as helical in Fig. 1 of the main text.

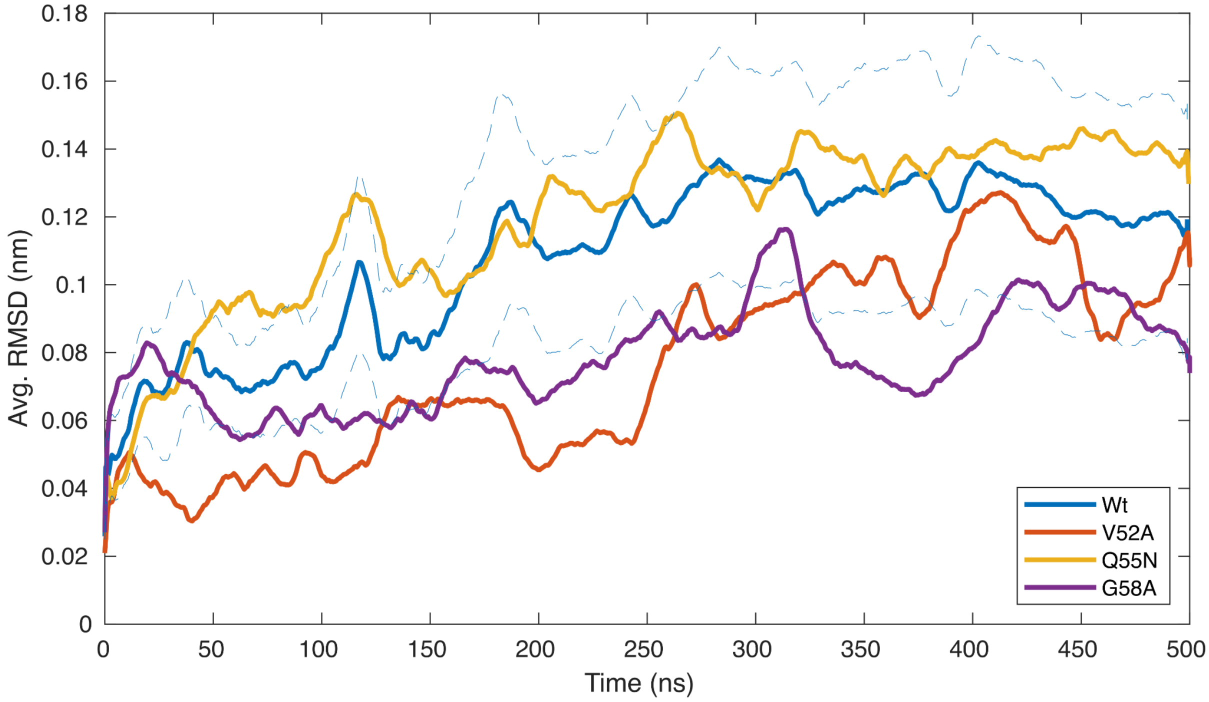


**Figure S4:** **Averages of the RMSD for all hinges for each LacI variant shown in Fig. S3.**

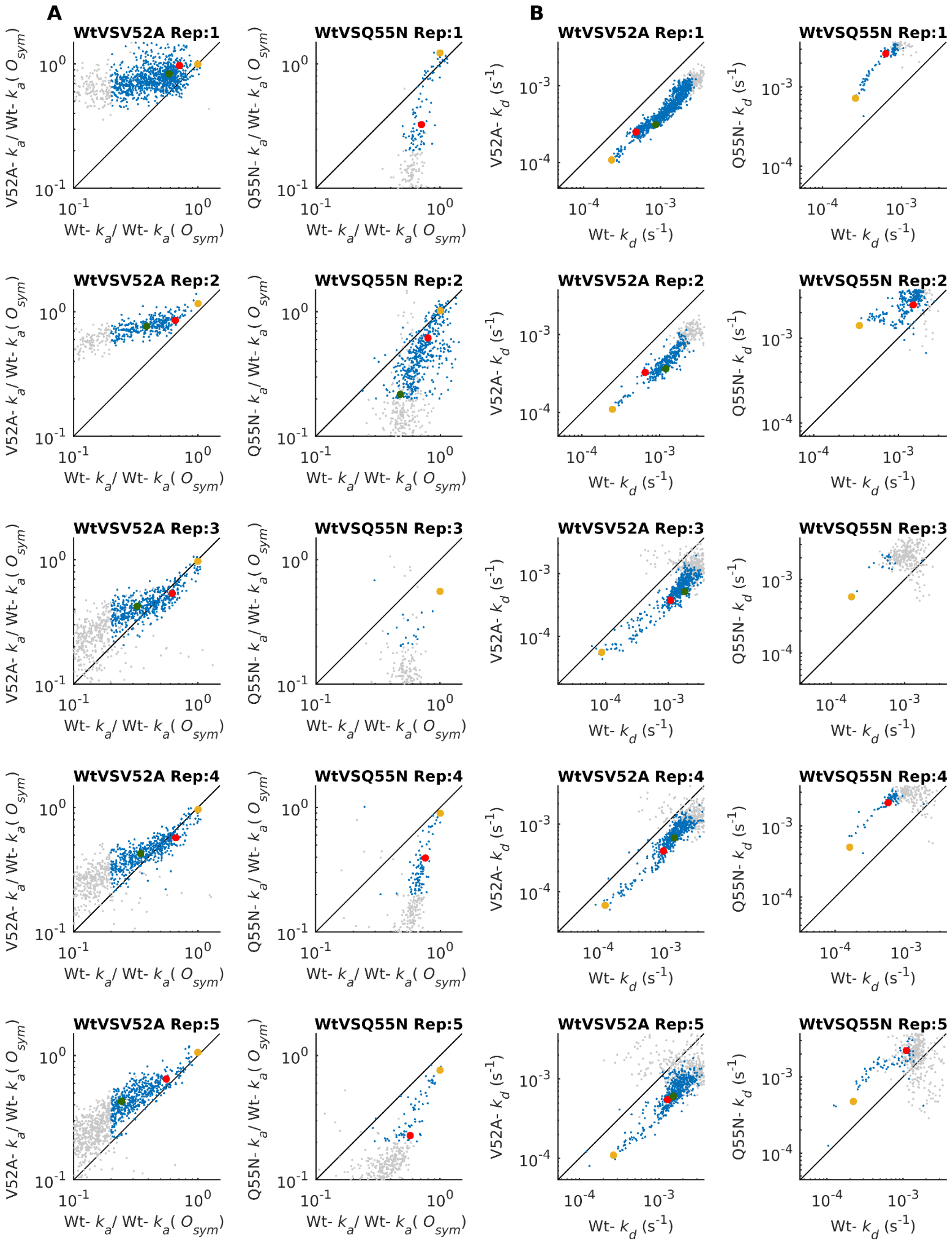
A
B
WtVSV52A Rep:1
WtVSQ55N Rep:1
WtVSV52A Rep:2
WtVSQ55N Rep:2
WtVSV52A Rep:3
WtVSQ55N Rep:3
WtVSV52A Rep:4
WtVSQ55N Rep:4
WtVSV52A Rep:5
WtVSQ55N Rep:5
V52A- $k_a$/ Wt- $k_a$( $O_{sym}$)
Q55N- $k_a$/ Wt- $k_a$( $O_{sym}$)
Wt- $k_a$/ Wt- $k_a$( $O_{sym}$)
V52A- $k_d$ ($s^{-1}$)
Q55N- $k_d$ ($s^{-1}$)
Wt- $k_d$ ($s^{-1}$)
$10^{-1}$
$10^{0}$
$10^{-4}$
$10^{-3}$

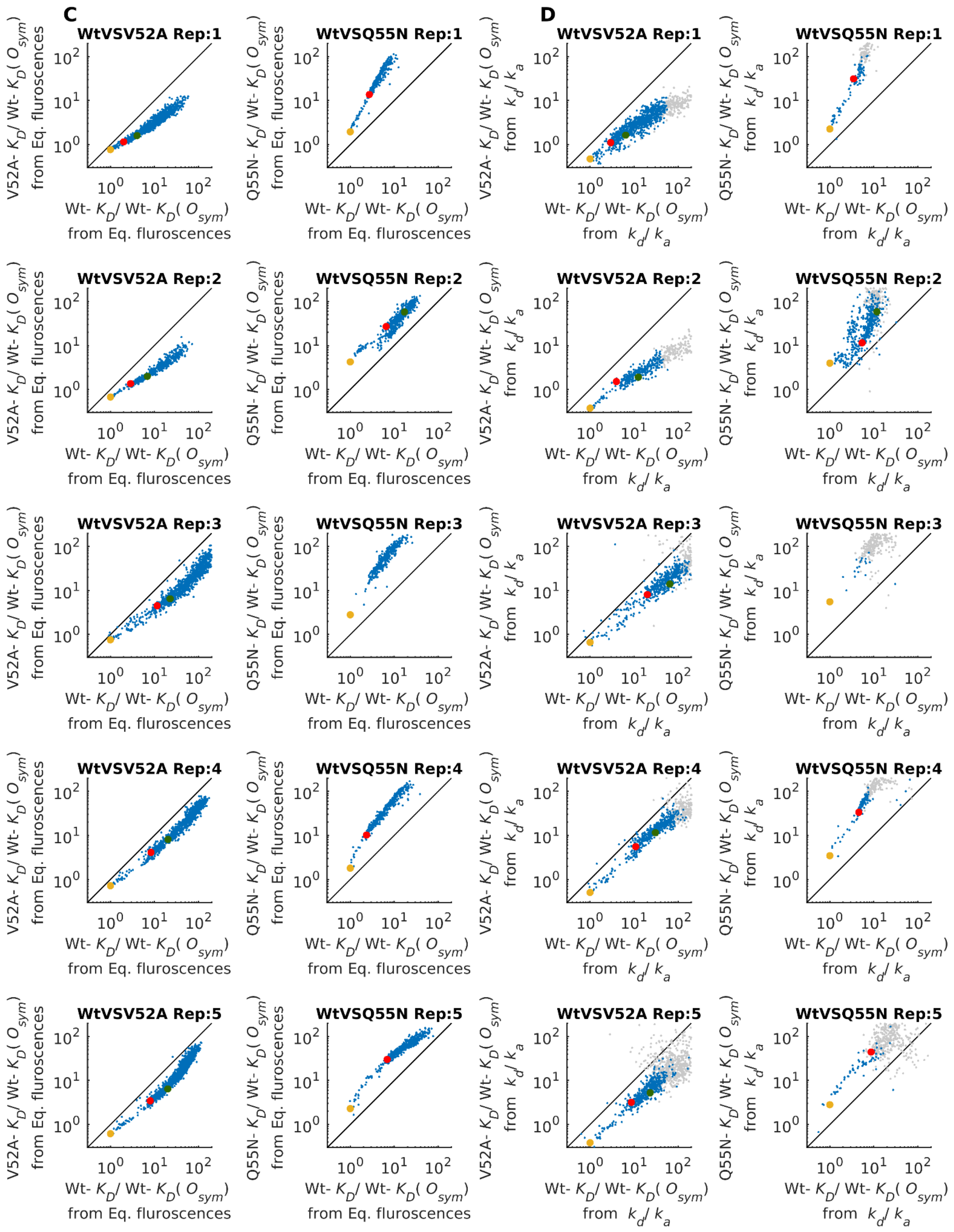


**Figure S5 Individual Wt VS Mutant side by side experimental replicates for data shown in Figure 2C, D&E.** In each replicate of the side-by-side PBM experiment, sequences with high $k_a$ and $k_d$ variability (CV > 1) or relative $k_a$ (normalized to Wt- $k_a$ ($O_{sym}$) within each replicate)<0.2 are shown in grey. $K_D$ values calculated from $k_a$ $k_d$ measurements meeting CV > 1 or relative $k_a$ <0.2 are also marked in grey.

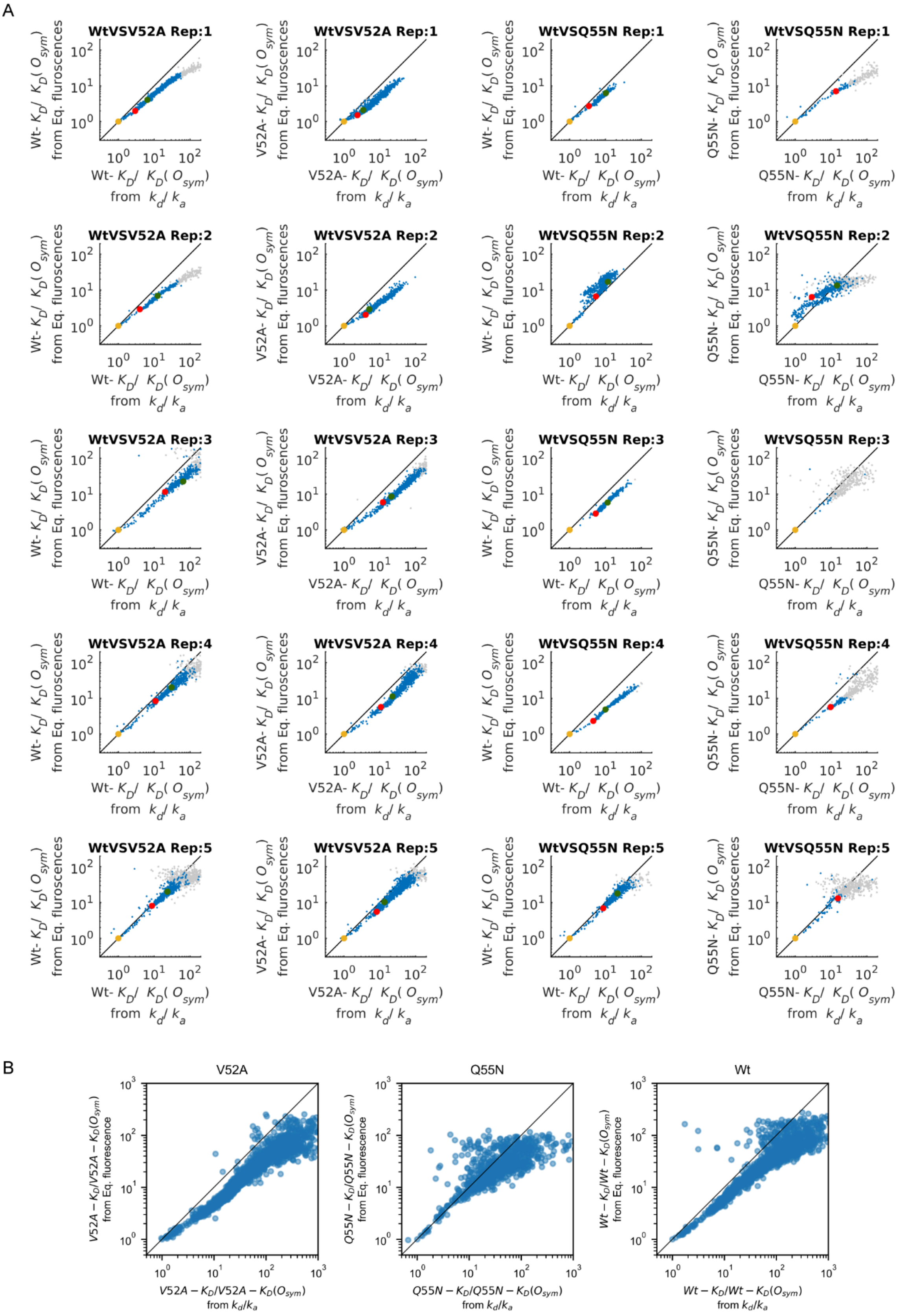


**Figure S6 Correlation of relative $K_d$ estimates from two different methods for calculating $K_d$. A.** Individual correlation of $K_d$ estimates from two methods for each protein in each experiment. **B.** Correlation of mean $K_d$ estimates from two methods, each data point of $K_d$ is taken from the mean value of 10 replicates for Wt, 5 replicates for V52A and 5 replicates for Q55N.

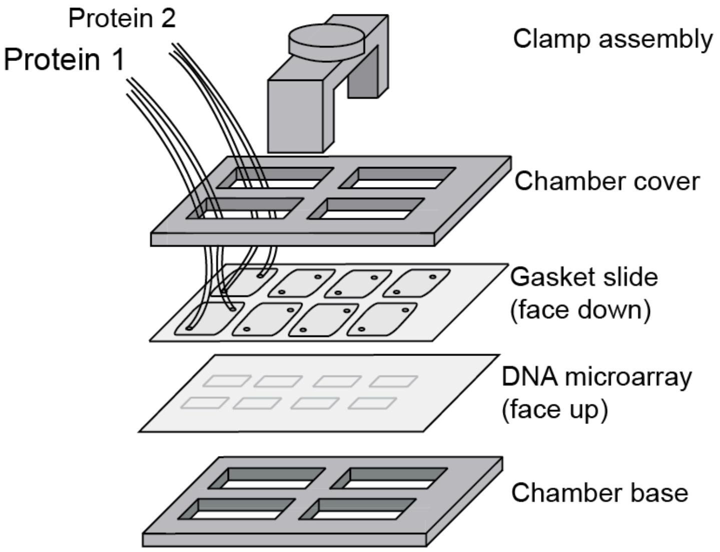


**Figure S7 Device set up for measuring kinetics in vitro with Protein binding Microarrays**

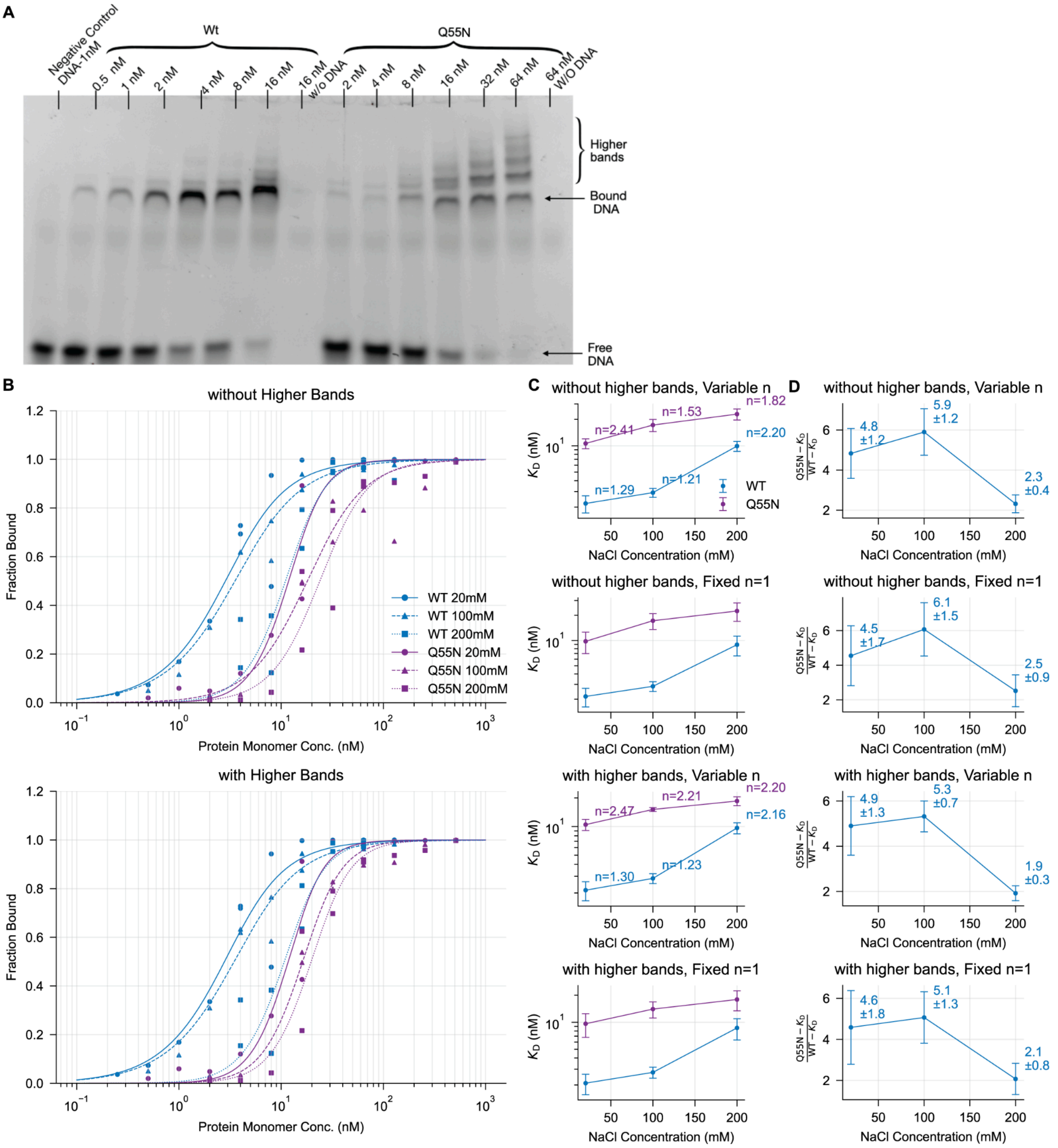


**Figure S8 EMSA-salt titration for Wt-LacI and Q55N-LacI** **A**. Example of Electrophoretic Mobility Shift Assay (EMSA) gel showing the binding of wild-type (WT) and Q55N mutant LacI to DNA (a 60bp DNA fragment including $O_{sym}$) at six different protein monomer concentrations in a 100 mM salt condition. The higher bands of DNA, LacI-bound DNA and free DNA are indicated in the image. note: LacI concentration labelled here are all for LacI monomers. **B**. Binding curves for the fraction of LacI-bound DNA were fitted using two models: a simple binding model with a fixed Hill coefficient (n=1) and a Hill model with a variable Hill coefficient (n). In the simple binding model, the fraction of LacI-bound DNA, f, is described by $f(L, Kd, D) = \frac{(L+D+Kd)-\sqrt{(L+D+Kd)^2-4LD}}{2D}$ calculates the fraction of DNA bound by solving the quadratic equation for the equilibrium between protein (L, denoted in equation for total protein concentration), DNA (D, denoted in equation for total DNA concentration ), and the dissociation constant (Kd). The Hill model, solves a system of two equations incorporating the Hill equation $\theta = \frac{Lfree^n}{Kd^n+Lfree^n}$ and the conservation of mass $L_{free} = L_{total} -$

*n*D*θ*, where $\theta$ is the fraction of DNA bound, $L_{free}$ is the free protein concentration, $L_{total}$ is the total protein concentration, and *n* is the Hill coefficient. The system is solved numerically for each protein tested to determine its *Kd* and Hill coefficient in three salt concentrations. The best-fit model was selected based on the lowest residual sum of squares and then is plotted. **C.** Equilibrium dissociation constants ($K_d$) determined from B for Wt-LacI and Q55N-LacI at varying NaCl concentrations. Data are shown as $K_d$ values with their corresponding standard errors, comparing salt-dependent inverse of affinities across four distinct analysis scenarios **D**. Corresponding ratios of Q55N-LacI Kd to Wt-LacI Kd at each salt concentration, illustrating relative changes in salt sensitivity. Error bars represent propagated standard errors from panel C.

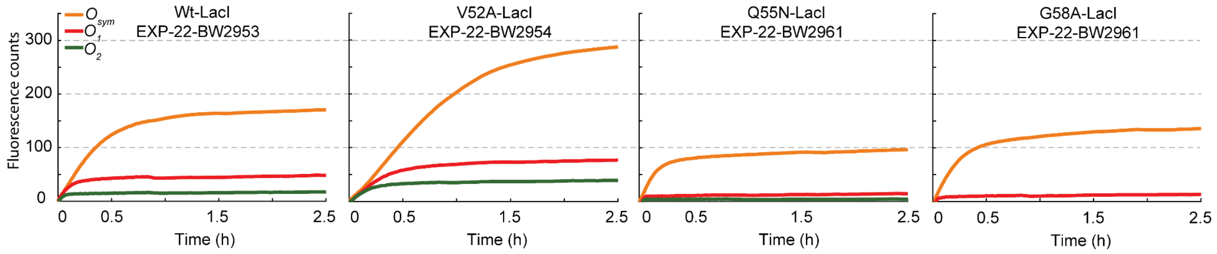


**Figure S9 Initial screening: Binding curves for four LacI variants (WT, V52A, G55N, G58A) to $O_{sym}$ (orange), $O_1$ (red), and $O_2$ (green) operators.** Plots display fluorescence counts over time with the negative control signal subtracted.

### Supplementary Tables:

**Table S1. Bacteria strains used in this study**

| Strains | Genotype or Relevant characteristics[1] | Reference or source |
|---|---|---|
| **Microscopy (single molecule experiment)** | | |
| EL8 | *Source for selectable/counterselectable marker, Acatsac1* | Gift from Dan I. Andersson lab (DA58376) |
| EL2620 | Δ*fliA*::Kan$^R$ *(Source for kanamycin* resistance cassette) | Jones et al., unpublished |
| EL3498 | *lacI-mVenus* Δ($p_{lac}$-*lacZ*)::term | This work |
| EL3518 | *lacI(V52A)-mVenus* Δ($p_{lac}$-*lacZ*)::term | This work |
| EL3520 | *lacI(Q55N)-mVenus* Δ($p_{lac}$-*lacZ*)::term | This work |
| EL4053 | *lacI-mVenus* Δ($p_{lac}$-*lacZ*)::term *aslA-glmZ* intergenic::cat- lacOsym | This work |
| EL4089 | *lacI-mVenus* Δ($p_{lac}$-*lacZ*)::term *aslA-glmZ* intergenic::cat-$O_{sym}$-*lacZ* | This work |
| EL4210 | Kan$^R$-$p_{FAB138}$-*lacI-mVenus* Δ($p_{lac}$-*lacZ*)::term | This work |
| EL4219 | Kan$^R$-$p_{FAB138}$-*lacI-mVenus* Δ($p_{lac}$-*lacZ*)::term | This work |
| EL4226 | Kan$^R$-$p_{FAB138}$-*lacI-venus* Δ($p_{lac}$-*lacZ*)::term aslA-glmZ intergenic::cat-lacOsym | This work |
| EL4242 | *lacI(V52A)-mVenus* Δ($p_{lac}$-*lacZ*)::term *aslA-glmZ* intergenic::cat- lacOsym | This work |
| EL4243 | *lacI(Q55N)-mVenus* Δ($p_{lac}$-*lacZ*)::term *aslA-glmZ* intergenic::cat- lacOsym | This work |
| EL4264 | Kan$^R$-$p_{FAB138}$-*lacI(V52A)-mVenus* Δ($p_{lac}$-*lacZ*)::term | This work |
| EL4265 | Kan$^R$-$p_{FAB138}$-*lacI(Q55N)-mVenus* Δ($p_{lac}$-*lacZ*)::term | This work |
| EL4266 | Kan$^R$-$p_{FAB138}$-*lacI(V52A)*-mVenus Δ($p_{lac}$-*lacZ*)::term | This work |
| EL4267 | Kan$^R$-$p_{FAB138}$-*lacI(Q55N)-mVenus* Δ($p_{lac}$-*lacZ*)::term | This work |

| | | |
|---|---|---|
| EL4268 | Kan$^R$-$p_{FAB138}$-*lacI(V52A)-mVenus* Δ($p_{lac}$-*lacZ*)::term *aslA-glmZ* intergenic::cat-lacOsym | This work |
| EL4269 | Kan$^R$-$p_{FAB138}$-*lacI(V52A)-mVenus* Δ($p_{lac}$-*lacZ*)::term *aslA-glmZ* intergenic::cat-lacOsym | This work |
| **Strains for Miller assay** | | |
| EL409 | *lacI-mVenus O1-* lacOsym *O2-* | This work |
| EL4491 | Kan$^R$-$p_{FAB138}$-*lacI-mVenus* Δ($p_{lac}$-*lacZ*)::term *aslA-glmZ* intergenic::cat-lacOsym-*lacZ* | This work |
| EL4493 | Kan$^R$-$p_{FAB138}$-*lacI(V52A)-mVenus* Δ($p_{lac}$-*lacZ*)::term *aslA-glmZ* intergenic::cat-lacOsym-*lacZ* | This work |
| EL4495 | Kan$^R$-$p_{FAB138}$-*lacI(Q55N)-mVenus* Δ($p_{lac}$-*lacZ*)::term *aslA-glmZ* intergenic::cat-lacOsym-*lacZ* | This work |
| EL4536 | Kan$^R$-$p_{FAB138}$-*mVenus Δ(lacI-lacZ)*::term | This work |
| EL4580 | Kan$^R$-$p_{FAB138}$-*mVenus Δ(lacI-lacZ)*::term *aslA-glmZ*::cat-lacOsym-*lacZ* | This work |
| EL4621 | *Δ(mhpR-lacZ)*::catsacB | This work |
| EL4630 | Kan$^R$-$p_{FAB138}$-*lacI-halo Δ(mhpR-lacZ)* | This work |
| EL4632 | Kan$^R$-$p_{FAB138}$-*lacI*(V52A)-*halo Δ(mhpR-lacZ)* | This work |
| EL4634 | Kan$^R$-$p_{FAB138}$-*lacI*(Q55N)-*halo Δ(mhpR-lacZ)* | This work |
| EL4639 | Kan$^R$-$p_{FAB138}$-*lacI-halo Δ(mhpR-lacZ) aslA-glmZ* intergenic::cat-lacOsym-*lacZ* | This work |
| EL4641 | Kan$^R$-$p_{FAB138}$-*lacI*(V52A)-*halo Δ(mhpR-lacZ)* *aslA-glmZ* intergenic::cat-lacOsym-*lacZ* | This work |
| EL4643 | Kan$^R$-$p_{FAB138}$-*lacI*(Q55N)-*halo Δ(mhpR-lacZ)* *aslA-glmZ* intergenic::cat-lacOsym-*lacZ* | This work |

[1]cat, gene encoding for CAT (Chloramphenicol acetyl-transferase)

**Table S2. Primers used in this study**

| Primers | Sequence (5'3')[1] | Comments |
|---|---|---|
| aslA-hom-cat-Osym -Fwd | aggagagcgttttcaatcctacctctg gcgcagttgatatTTACGCCCC GCCCTGCCAC | Primer to obtain fragment cat-lacOsym or cat-lacO$_{sym}$-lacZ fragment; Forward primer |
| aslA-hom-cat-Osym -Rev | agcatcaataatcaacgcgatataata aacctgccttacaattgtgagcgctca caattCACGTAAGAGGTTC CAACTTTCACC | Primer to obtain cat-lacOsym fragment; Reverse primer |
| cat-Osym-lacZ_aslA _R | agcatcaataatcaacgcgatataata aacctgccttacTTATTTTTGA CACCAGACCAACTGG | Primer to obtain cat-lacO$_{sym}$-lacZ fragment; Reverse primer |
| Comp2_lacI 14 bp_RBS_p138-Fw | gttactggtttcacattcaccaccctga attgactctcttatacgttaaatctacga gccggatgattaattgtctaacaccgt gCGCAAGCGCAAAGAG AAAGC | Primer to obtain Kan$^{R}$-pFAB138-RBS-lacI(14bp) fragment; Forward primer |
| Comp1_Kan-Rv | gtgcaccaggtgcaccacgttgtttta actatagaaatgTCAGAAGAA CTCGTCAAGAAGG | Primer to obtain Kan$^{R}$-pFAB138-RBS-lacI(14bp) fragment; Reverse primer |

| | | |
|---|---|---|
| LacI_Halo_R | cctattctctagaaagtatagggcaattctcgagcacaagTTATTAACCGTGATGGTGATGG | Primer to obtain fragments, KanR-pFAB138-lacI-halo or KanR-pFAB138-lacI(V52A)-halo or KanR-pFAB138-lacI(Q55N)-halo; Reverse |
| insert aslA-glmZ_fw | GATAATTGAGATCCCTCTCCC | Sequencing primers to verify insert at aslA-glmZ intergenic region |
| insert aslA-glmZ_rv | ACCTGAGCTTGATCCTACAC | Sequencing primers to verify insert at aslA-glmZ intergenic region |
| lacI insert_mhpA_fw | GGTTAACAGCAGGCTGGATGTC | Sequencing primer to verify *lacI-mVenus* |
| kan_insert_seq | CAACCTTACCAGAGGGCGCC | Sequencing primer to verify KanR-pFAB138-*lacI-mVenus* |
| venus-lacY-DN-conf - rv | CCACAGCAGGTATTTGCGCAGC | Sequencing primer to verify *lacI-mVenus* |
| CAT-R | GCAACTGACTGAAATGCCTC | Sequencing primer to verify cat-lacOsym or cat-lac$O_{sym}$-lacZ insert |
| T7 Term_fwd | CTAGCATAACCCCTTGGGGC | Sequencing primer to verify *lacI-halo* insert |
| Seq_Halo_F | AGTTCATCCGCCCTATCCC | Sequencing primer to verify *lacI-halo* insert |
| Seq_Halo_R | CTTCGACCAGCGCGACGATG | Sequencing primer to verify *lacI-halo* insert |

| | | |
|---|---|---|
| lacI mid rev3252 | ACGCGCCGAGACAGAA CTTA | Sequencing primer *lacI-halo* insert |

[1]Nucleotide sequences in lowercase are the 40 bases homology for lambda red

**Table S3. Sequences of LacI-Halo variants for Protein binding Microarray assay**

The Halotag used for fluorescent labelling, which was introduced in the C terminal of the LacI sequence is underlined. The linker sequence between the Halotag and the original LacI sequence is marked in bold. Mutations for LacI are marked in color. 6x-His tags were introduced for protein purification.

| Name | Sequence |
|---|---|
| LacI-Halo | MKPVTLYDVAEYAGVSYQTVSRVVNQASHVSAKTREKVEAAMAEL<br>NYIPNRVAQQLAGKQSLLIGVATSSLALHAPSQIVAAIKSRADQLGAS<br>VVVSMVERSGVEAAKAAVHNLLAQRVSGLIINYPLDDQDAIAVEAA<br>ATNVPALFLDVSDQTPINSIIFSHEDGTRLGVEHLVALGHQQIALLAGP<br>LSSVSARLRLAGWHKYLTRNQIQPIAEREGDWSAMSGFQQTMQML<br>NEGIVPTAMLVANDQMALGAMRAITESGLRVGADISVVGYDDTEDS<br>SCYIPPLTTIKQDFRLLGQTSVDRLLQLSQGQCVKGNQLLPVSLVKRK<br>TTLAPNTQT**G**AEIGTGFPFDPHYVEVLGERMHYVDVGPRDGTPVLF<br>LHGNPTSSYVWRNIIPHVAPTHRCIAPDLIGMGKSDKPDLGYFFDDH<br>VRFMDAFIEALGLEEVVLVIHDWGSALGFHWAKRNPERVKGIAFME<br>FIRPIPTWDEWPEFARETFQAFRTTDVGRKLIIDQNVFIEGTLPMGVV<br>RPLTEVEMDHYREPFLNPVDREPLWRFPNELPIAGEPANIVALVEEYM<br>DWLHQSPVPKLLFWGTPGVLIPPAEAARLAKSLPNCKAVDIGPGLNL<br>LQEDNPDLIGSEIARWLSTLEISGHHHHHHG |
| V52ALacI-Halo | MKPVTLYDVAEYAGVSYQTVSRVVNQASHVSAKTREKVEAAMAEL<br>NYIPNRAAQQLAGKQSLLIGVATSSLALHAPSQIVAAIKSRADQLGAS<br>VVVSMVERSGVEAAKAAVHNLLAQRVSGLIINYPLDDQDAIAVEAA<br>ATNVPALFLDVSDQTPINSIIFSHEDGTRLGVEHLVALGHQQIALLAGP<br>LSSVSARLRLAGWHKYLTRNQIQPIAEREGDWSAMSGFQQTMQML<br>NEGIVPTAMLVANDQMALGAMRAITESGLRVGADISVVGYDDTEDS<br>SCYIPPLTTIKQDFRLLGQTSVDRLLQLSQGQCVKGNQLLPVSLVKRK<br>TTLAPNTQT**G**AEIGTGFPFDPHYVEVLGERMHYVDVGPRDGTPVLF<br>LHGNPTSSYVWRNIIPHVAPTHRCIAPDLIGMGKSDKPDLGYFFDDH<br>VRFMDAFIEALGLEEVVLVIHDWGSALGFHWAKRNPERVKGIAFME<br>FIRPIPTWDEWPEFARETFQAFRTTDVGRKLIIDQNVFIEGTLPMGVV<br>RPLTEVEMDHYREPFLNPVDREPLWRFPNELPIAGEPANIVALVEEYM<br>DWLHQSPVPKLLFWGTPGVLIPPAEAARLAKSLPNCKAVDIGPGLNL<br>LQEDNPDLIGSEIARWLSTLEISGHHHHHHG |
| Q55NLacI-Halo | MKPVTLYDVAEYAGVSYQTVSRVVNQASHVSAKTREKVEAAMAEL<br>NYIPNRVAQNLAGKQSLLIGVATSSLALHAPSQIVAAIKSRADQLGAS<br>VVVSMVERSGVEAAKAAVHNLLAQRVSGLIINYPLDDQDAIAVEAA<br>ATNVPALFLDVSDQTPINSIIFSHEDGTRLGVEHLVALGHQQIALLAGP<br>LSSVSARLRLAGWHKYLTRNQIQPIAEREGDWSAMSGFQQTMQML<br>NEGIVPTAMLVANDQMALGAMRAITESGLRVGADISVVGYDDTEDS<br>SCYIPPLTTIKQDFRLLGQTSVDRLLQLSQGQCVKGNQLLPVSLVKRK<br>TTLAPNTQT**G**AEIGTGFPFDPHYVEVLGERMHYVDVGPRDGTPVLF<br>LHGNPTSSYVWRNIIPHVAPTHRCIAPDLIGMGKSDKPDLGYFFDDH<br>VRFMDAFIEALGLEEVVLVIHDWGSALGFHWAKRNPERVKGIAFME<br>FIRPIPTWDEWPEFARETFQAFRTTDVGRKLIIDQNVFIEGTLPMGVV<br>RPLTEVEMDHYREPFLNPVDREPLWRFPNELPIAGEPANIVALVEEYM |

| | |
|---|---|
| | DWLHQSPVPKLLFWGTPGVLIPPAEAARLAKSLPNCKAVDIGPGLNL LQEDNPDLIGSEIARWLSTLEISGHHHHHHG |

**Table S4. Occurrences of Operator Binding Sites in the E. coli Genome**

The table lists the counts of operator mutation types located within the E. coli chromosome, with their relative positions to *OriC* indicated in brackets for mutations occurring fewer than 3 times.

| Mutation Category | Osym | O1 | O2 | Total |
|---|---|---|---|---|
| **No mutation** | **1( -61071)** | **0** | **0** | **1** |
| **Single Mutation** | **0** | **0** | **0** | **0** |
| **Double Mutations** | **0** | **0** | **0** | **0** |
| **Triple mutations** | **0** | **1 (+820407)** | **1 (+2844762)** | **4** |
| **Quadruple mutations** | **14** | **12** | **14** | **66** |
| **Total** | **15** | **13** | **15** | **71** |

**Table S5. Comparative parameters of Wt-LacI and hinge-helix mutants (V52A, Q55N) from *in vitro* and *in vivo* assays**

| Expriment Type | LacO Type | LacI-Halo variant | $k_a$/ Wt-$k_a(O_{sym})$ | $k_d$ (±SEM) x$10^{-4}$ $s^{-1}$ | $K_D$/ Wt-$K_D(O_{sym})$ | | | |
|---|---|---|---|---|---|---|---|---|
| ***In Vitro*: HT Kinetics Measurement on PBM** | $O_{sym}$ | **Wt** | 1 (ref.) | 2.14 ± 0.03 | 1 (ref.) | | | |
| | | **V52A** | ~1 | 0.89 ± 0.02 | ~0.4 (affinity ↑) | | | |
| | | **Q55N** | ~0.9 | 7.39 ± 0.13 | ~3.6 (affinity ↓) | | | |
| | $O_1$ | **Wt** | ~0.7 | 8.84 ± 0.12 | ~6.1 | | | |
| | | **V52A** | ~0.7 | 3.80 ± 0.06 | ~2.6 | | | |
| | | **Q55N** | ~0.4 | 22.79 ± 0.89 | ~28.4 | | | |
| | $O_2$ | **Wt** | ~0.4 | 13.25 ± 0.35 | ~14.1 | | | |
| | | **V52A** | ~0.6 | 4.84 ± 0.10 | ~4.1 | | | |
| | | **Q55N** | ~0.1 | 36.53 ± 3.80 | ~140.5 | | | |
| / | / | **LacI-mVenus Variant** | / | / | **- IPTG** | | **+ IPTG** | |
| | | | | | $\phi^1$ | $p_{bound}^2$ | $\phi^1$ | $p_{bound}^2$ |
| ***In vivo*: Miller assay** | $O_{sym}$ | **Wt** | / | / | ~38.9 | ~97% | ~1.3 | ~23% |
| | | **V52A** | / | / | ~57.7 | ~98% | ~8.8 | ~89% |
| | | **Q55N** | / | / | ~4.2 | ~76% | ~1 | ~0% |
| | | **ΔLacI** | / | / | 1 (ref.) | / | 1 (ref.) | / |
| / | | | $k_a^3$ $min^{-1}$ | / | **Δavg.N(dots)/cell[4]:** | | **Δavg.N(dots)/cell[4]:** | |
| ***In vivo*: Single-molecule assay** | $O_{sym}$ | **Wt** | ~1.3 | / | 0.34 ± 0.02 | | 0.04 ± 0.01 | |
| | | **V52A** | / | / | 0.28 ± 0.02 | | 0.33 ± 0.02 | |
| | | **Q55N** | ~1.3 | / | 0.20 ± 0.01 | | 0.00 ± 0.00 | |

[1] ***ϕ***: Repression strength, denoted detaily in the main text.

[2] $p_{bound}$: $O_{sym}$ occupancy, see derivation in **Methods**.

[3] $k_a$: Fitted $k_a$ values, taken from Figure 4C

[4]**Δavg.N(dots)/cell**: Taken from Figure 3E.

**Table S6: Lower bounds of distance restraints between alpha carbons that were used in MD simulation.**

| DBD on Chain A | | DBD on Chain B | |
|---|---|---|---|
| Residue pair | Lower bound (nm) | Residue pair | Lower bound(nm) |
| Thr34-Asp88 | 1.5 | Thr34-Asp88 | 2.0 |
| Thr34-Ala106 | 1.8 | Thr34-Ala106 | 2.2 |
| Thr34-Ala110 | 2.0 | Thr34-Ala110 | 2.0 |
| Glu39-Asp88 | 1.4 | Glu39-Asp88 | 1.5 |
| Glu39-Ala106 | 1.4 | Glu39-Ala106 | 1.5 |
| Glu39-Ala110 | 1.3 | Glu39-Ala110 | 1.8 |
| Ala43-Ala106 | 1.3 | Ala43-Ala106 | 1.4 |
| Ala43-Ala110 | 1.3 | Ala43-Ala110 | 1.3 |
